# GAPIN: Grouped and Aligned Protein Interface Networks

**DOI:** 10.1101/520833

**Authors:** Biharck M. Araújo, Aline L. Coelho, Sabrina A. Silveira, João P. R. Romanelli, Raquel C. de Melo-Minardi, Carlos H. da Silveira

## Abstract

**Summary:** GAPIN is a web-based application for structural interaction network analysis among any type of PDB molecules, regardless of whether their interfaces are between chain-chain or chain-ligand. A special emphasis is given to graph clustering, allowing users to scrutinize target contexts for ligand candidates. We show how GAPIN can be used to unveil underlying hydrophobic patterns on a set of peptidase-inhibitor complexes. In another experiment, we show there is a positive correlation between cluster sizes and the presence of druggable spots, indicating that the clustering may discriminate the higher complexity of these hot subnetworks.

**Availability and implementation:** GAPIN is freely available as an easy-to-use web interface at https://gapin.unifei.edu.br.

**Supplementary information:** Supplementary data are available online.

## Introduction

Several computational methods, databases, and original tools have been proposed to help analyze the profusion of biomolecular interaction data generated by experimentalists, hoping to extract useful knowledge mainly through comprehensive mappings (Kovács et al., 2019), such as NetworkAnalyzer/RINalyzer (Doncheva et al.,2012), STRING (Szklarczyk et al, 2017), Bio3D-web (Skjærven et al., 2016), NAPS (Chakrabarty et al.,2016), STITCH (Szklarczyk et al., 2016), among others (McGillivray et al.,2018). However, these initiatives usually have their own focus and they do not easily integrate different types of interfaces, such as chain-chain or chain-ligand, regardless of the biomolecule involved. Moreover, how can we best visualize these interaction networks with the design of new drug candidates in mind?

To join efforts to this challenge, we propose GAPIN - *Grouped and Aligned Protein Interface Networks*, a fully web-based application with immediate availability, portability, and all friendly convenience that a modern browser can offer. GAPIN has RCSB PDB (Berman et al., 2000) coordinates as main input, defining interfaces among biomolecules as graphs with node representations at the atomic level. This permits visualizations and analysis regardless of whether the interfaces are among proteins, nucleic acids, carbohydrates, lipids, chemical ligands, ions or even waters. GAPIN contrasts the PDB structures and correspondent graph models with different view options. It introduces a network modularity analysis, by clustering the atomic graphs into communities of densely connected nodes, forming higher-level or modular graphs. GAPIN provides a graph-structure alignment tool for comparative interface analysis. Higher-level graphs also help identify spots, a group of a few residues in chain-chain interactions that can have a relevant contribution to free energy of binding. Consequently, they can be candidates for new druggable targets (Xia et al., 2010). To highlight the wide scope of GAPIN, we perform two experiments: one on graph alignments in a peptidase-inhibitor data set (Gonçalves-Almeida et al., 2012), and another on spot locations in an alanine mutagenesis data set (Xia et al., 2010).

## Material and Methods

See Supplementary Material.

## Interface Alignments

Gonçalves-Almeida et al. (2012) showed how hydrophobic patterns in peptidase-inhibitor interfaces can help explain the cross-inhibition phenomenon, when dissimilar structural chains bind to a particular site promiscuously. Classical examples are some trypsin-like and subtilisin-like serine-peptidases. Notwithstanding their divergent 3D structures (all beta against alpha and beta) and sequence identity as low as 20%, they can be cross-inhibited by both Turkey ovomucoid or Eglin C, also two different inhibitors. The authors indicated the existence of conserved hydrophobic patches on such interfaces using a nonredundant set of 9 peptidase-inhibitor complexes. (see Table S2). Figure 1A shows the result of chain-chain interface alignments for 2 of these complexes, comprising nonpolar-nonpolar contacts in 1PPF (Leukocyte elastase - Turkey ovomucoid 3) and 1TEC (Thermitase subtilisin - Eglin C). It is possible to see the graph similarities and the notable superimposition of nodes, in agreement with aforementioned results, but with the differential that GAPIN performs it for any PDB chain-chain interfaces (chain-ligand interfaces soon). See Figure S2 and S3 for the remaining alignments and RMSD statistics, respectively.

**Fig. 1.**
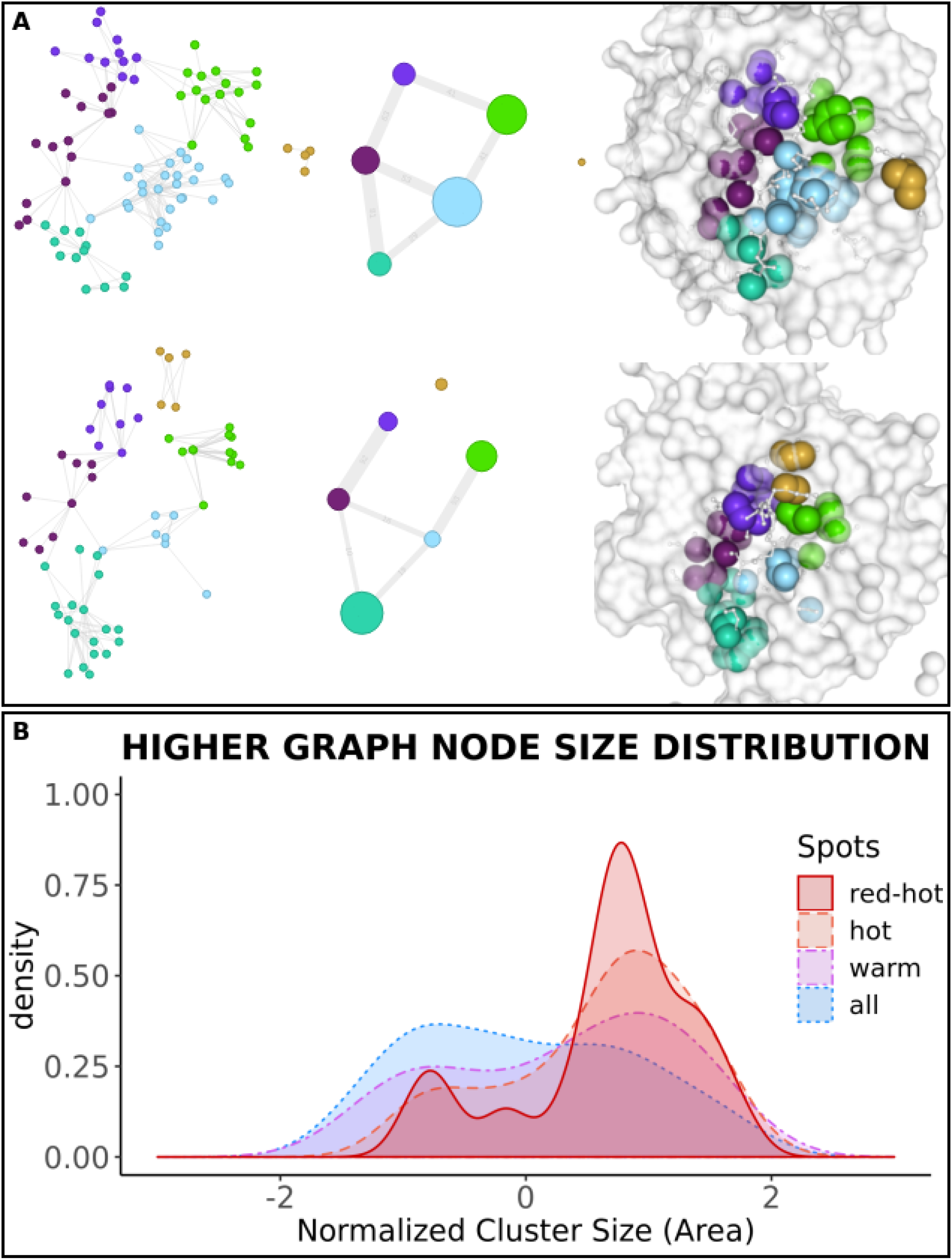
(**A**) Alignment results of nonpolar interfaces from two distinct serine-peptidase in complex with also two different inhibitors (PDB ids: 1PPF and 1TEC). (**B**) Node size distribution density in function of normalized cluster sizes (z-scores of the contact areas, using a higher graph minimum quality of 0.80).

## Spot Locations

At chain-chain interfaces, alanine mutations are the common procedure used to discriminate residues that account for most of the free energy of binding (Xia et al., 2010). If the impact of this mutation causes a change in ΔG of binding at least 2.0 kcal.mol^−1^, then the target residue is called a hot spot. If less than this, a warm spot. If greater than 4.0 kcal.mol^−1^, a red-hot spot. Xia et al. compiled an experimentally determined spot data set comprising 15 protein-protein complexes (see “Additional file 1” in this citation). We mapped the presence of these spots into the higher graphs, according to the node location. Node size distribution plots can be seen in Figure 1B for red-hot spots (red/solid), hot spot (orange/dashed) and warm spots (purple/dotdash) against the others (blue/dotted). The difference between the distributions is statically significant (at confidence level of 0.95) for red-hot and hot spots (KS-test with p-value 0.012 and 0.019, respectively) but not for warm spots (p-value 0.32). This indicates that larger nodes may be more likely to host druggable spots. GAPIN allows users to isolate and explore such nodes in different ways (Figure S4). See Table S3 for more statistical evidence.

## Conclusion

GAPIN is a modern web-based application focused on structural interaction network analysis of PDB biomolecule interfaces. Herein, we show how it can be used to unveil hydrophobic patterns between different peptidase-inhibitor complexes. We also show there is a positive correlation between node sizes and the presence of energetic spots. Probably because these interface regions involve more complex shape complementarity and denser interactions, resulting in more intricate subnetworks, which is in line with other studies (Moreira et. al.,2007).

## Supporting information

Tutorial material

GAPIN_supporting_raw_data_files-v2

## Acknowledgements

We are grateful to Prof. Ricardo Leão/USP by the inspiration of the name GAPIN.

