## Supplementary material for "GAPIN: Grouped and Aligned Protein Interface Networks": Tutorial material

[1] Bioinformatics PhD Program, Universidade Federal de Minas Gerais, 31270-901, Brazil.

[2] School of Medicine - EMED / Universidade Federal de Ouro Preto, 35400-000, Brazil.

[3] Department of Computer Science, Universidade Federal de Viçosa, 36570-900, Brazil.

[4] European Molecular Biology Laboratory, European Bioinformatics Institute, Wellcome Genome Campus, Hinxton, UK.

[5] Institute of Applied and Pure Sciences, Universidade Federal de Itajubá, 35903-087, Brazil.

[6] Department of Computer Science, Universidade Federal de Minas Gerais, 31270-901, Brazil.

[7] Institute of Technological Sciences, Universidade Federal de Itajubá, 35903-087, Brazil.


### Introduction

This tutorial section will walk you through about GAPIN's implementation, how to use it to get the best of analysis such as Figure S1 and present a study case in the end of it.

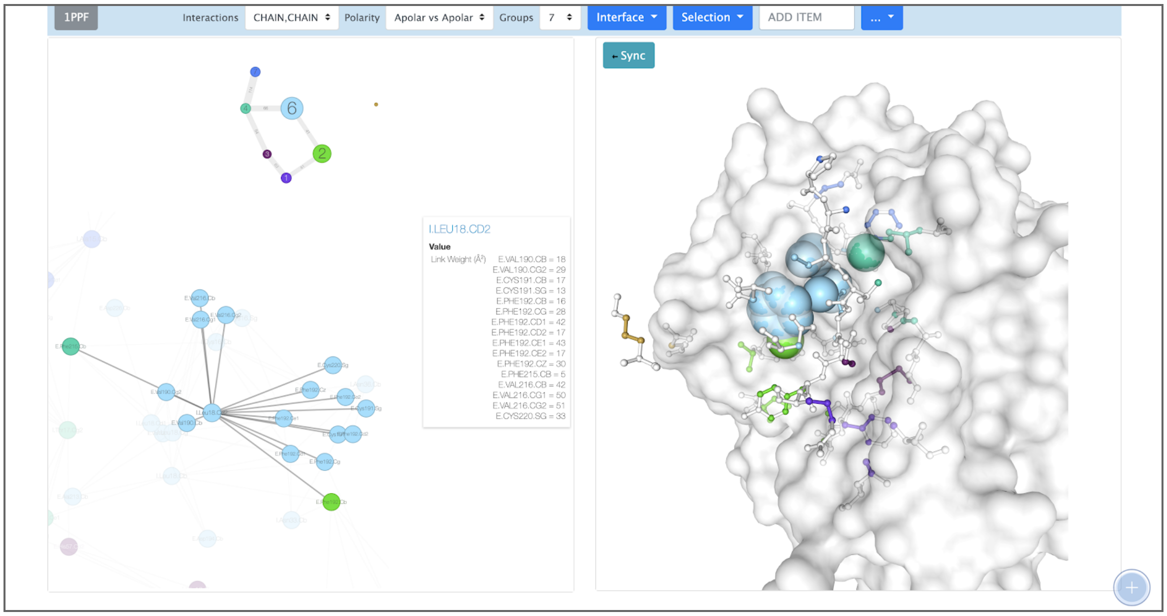


**Figure S1.** Analyzing the PDB 1PPF (X-ray crystal structure of the complex of human leukocyte elastase 2 (pmn elastase) and the third domain of the turkey ovomucoid inhibitor). Where can seem the first level interaction of the Beta Carbon from Leucine 18 from the Inhibitor.

### Implementation details

#### Architecture

GAPIN was built following Continuous Integration (Duvall et al., 2007) and Continuous Delivery (Humble, J., 2010) practices, Figure S2, where every time a new version has a green build, it is immediately deployed to the production environment. GAPIN's code base is currently stored at Bitbucket (Bitbucket, 2012) and using Bitbucket pipelines (Bitbucket Pipelines, 2019) for the Continuous Delivery and deployment process.

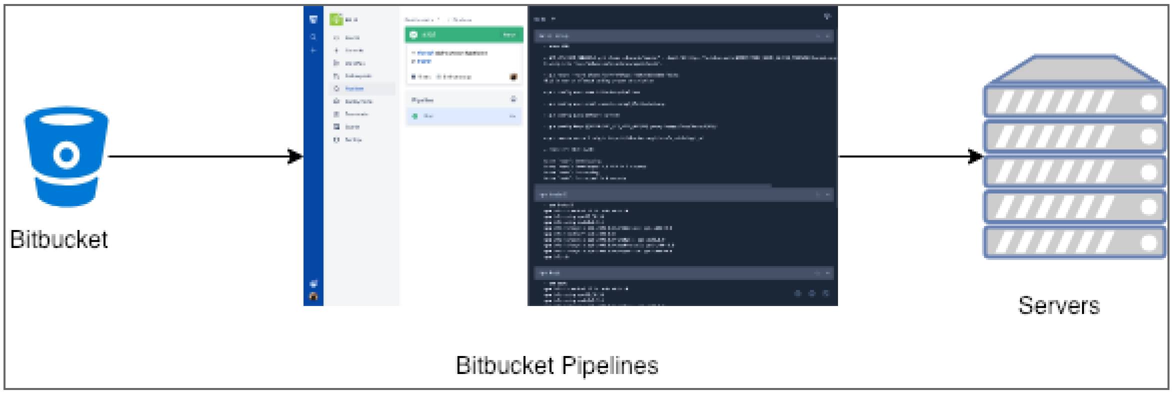


**Figure S2.** Version Control and Continuous Delivery process with Bitbucket and Bitbucket Pipelines

#### Back end

GAPIN's back end architecture is based on an asynchronous pattern with NodeJS (Casciaro, Mario., 2016) where process run in the background and generating promises

(Gallaba, K., Mesbah, A., Beschastnikh, I., 2015). Basically as Figure S3 shows, the back end part of GAPIN is divided into 3 parts: 1) GAPIN's back end application built in NodeJS (NodeJS, 2019) and ExpressJS (ExpressJS, 2019); 2) GAPIN's back end processor written in R (R Core Team., 2018); 3) GAPIN's storage with MongoDB (MongoDB, 2019).

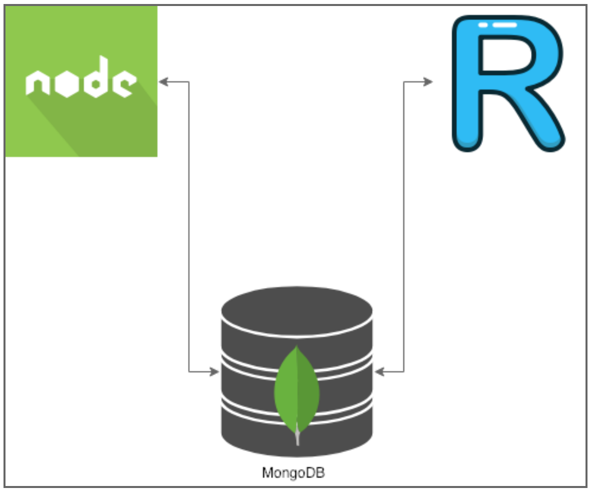


Figure S3. GAPIN's Back-end architecture

Based on that architecture, GAPIN takes advantage of MongoDB which stores documents used to load the protein on the front end part without any processing step, making the user experience good due to the velocity to load any data, such as the atoms that belong to the interface network, Table S1.


```
{
  "pdb": "1TEC",
  "pdb_type": 1,
  "title": "CRYSTALLOGRAPHIC REFINEMENT BY INCORPORATION OF MOLECULAR 2 DYNAMICS. THE
THERMOSTABLE SERINE PROTEASE THERMITASE 3 COMPLEXED WITH EGLIN-C",
  "bsr": [
    {
      "source": "E.SER107.O",
      "source_polarity": "POLAR",
```

```
      "target": "I.TYR35.CB",
      "target_polarity": "NONPOLAR",
      "bsr_value": 6.9
    }
  ]
}
```


**Table S1**. JSON sample stored at MongoDB as part of the GAPIN's process

#### Front end

GAPIN was built focused on simplicity, where users could interact with the system smoothly, grab pieces of information and discover easily. The styling layout was built with Bootstrap 4 (Otto, M., & Thornton, J., 2015), which by default is responsive enabling users to access the application from different devices such as Laptops, Tablets, Smartphones and so on.

The front end framework used by GAPIN is Handlebars (HandleBars JS., 2019) and the information which feed the front end comes from NodeJS, onto the back end part. The interactive part was written in NGL (Rose, A. S., *et al*., 2018) for macromolecules interactions, D3Plus (D3plus, 2018) for contact network connections and DataTables (DataTables, 2008) for interactive and searchable Jobs information.

#### Data availability and interoperability

GAPIN allows the user to download a CSV file based on PDB (Kouranov, A., *et al.,* 2005) format that users get the PDB adjacency matrix in relation to the interactions (Chain by Chain or Any by Ligand) and polarity (Polar by Polar, Apolar by Apolar or them all).

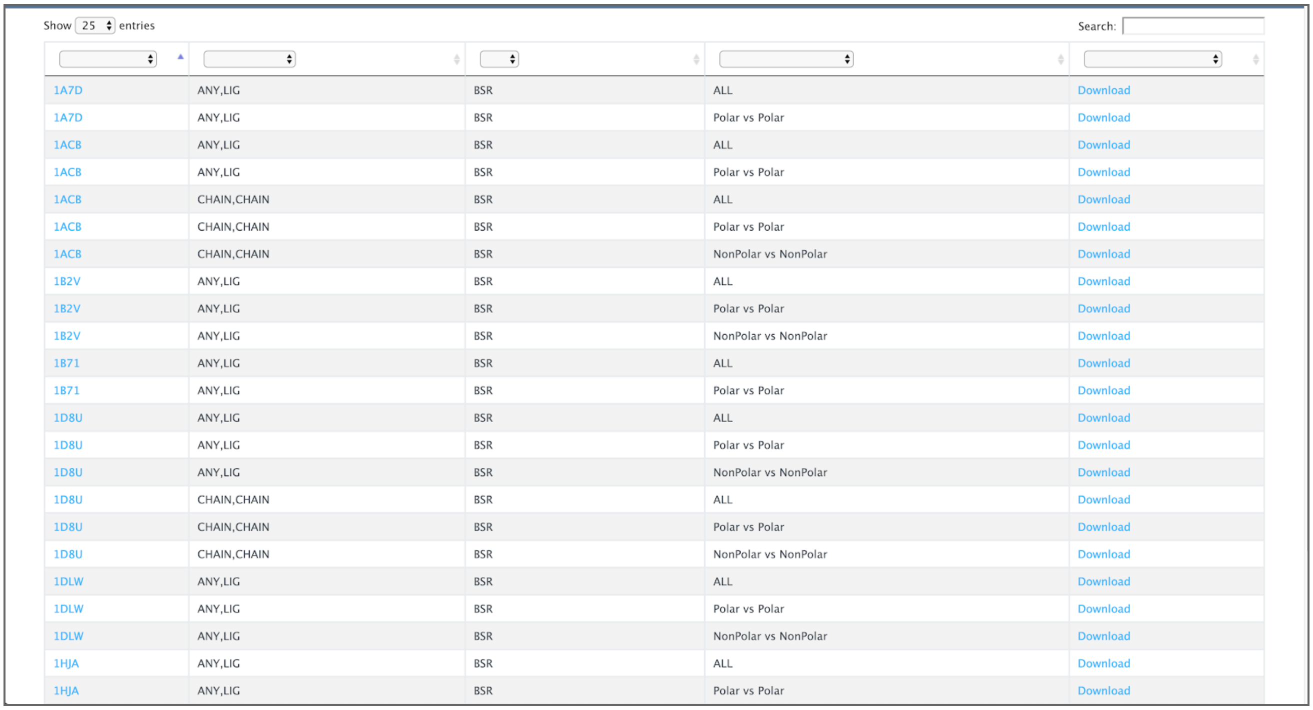

Show 25 entries | Search:

| | | | | |
|---|---|---|---|---|
| 1A7D | ANY,LIG | BSR | ALL | Download |
| 1A7D | ANY,LIG | BSR | Polar vs Polar | Download |
| 1ACB | ANY,LIG | BSR | ALL | Download |
| 1ACB | ANY,LIG | BSR | Polar vs Polar | Download |
| 1ACB | CHAIN,CHAIN | BSR | ALL | Download |
| 1ACB | CHAIN,CHAIN | BSR | Polar vs Polar | Download |
| 1ACB | CHAIN,CHAIN | BSR | NonPolar vs NonPolar | Download |
| 1B2V | ANY,LIG | BSR | ALL | Download |
| 1B2V | ANY,LIG | BSR | Polar vs Polar | Download |
| 1B2V | ANY,LIG | BSR | NonPolar vs NonPolar | Download |
| 1B71 | ANY,LIG | BSR | ALL | Download |
| 1B71 | ANY,LIG | BSR | Polar vs Polar | Download |
| 1D8U | ANY,LIG | BSR | ALL | Download |
| 1D8U | ANY,LIG | BSR | Polar vs Polar | Download |
| 1D8U | ANY,LIG | BSR | NonPolar vs NonPolar | Download |
| 1D8U | CHAIN,CHAIN | BSR | ALL | Download |
| 1D8U | CHAIN,CHAIN | BSR | Polar vs Polar | Download |
| 1D8U | CHAIN,CHAIN | BSR | NonPolar vs NonPolar | Download |
| 1DLW | ANY,LIG | BSR | ALL | Download |
| 1DLW | ANY,LIG | BSR | Polar vs Polar | Download |
| 1DLW | ANY,LIG | BSR | NonPolar vs NonPolar | Download |
| 1HJA | ANY,LIG | BSR | ALL | Download |
| 1HJA | ANY,LIG | BSR | Polar vs Polar | Download |

**Figure S3**. The export section where users can download the output from the atom interactions.

Throughout the interaction with GAPIN, users can also take and download unlimited screenshots, Figure S4, any time they desire for the macromolecule structure, atom contact network and groups network.

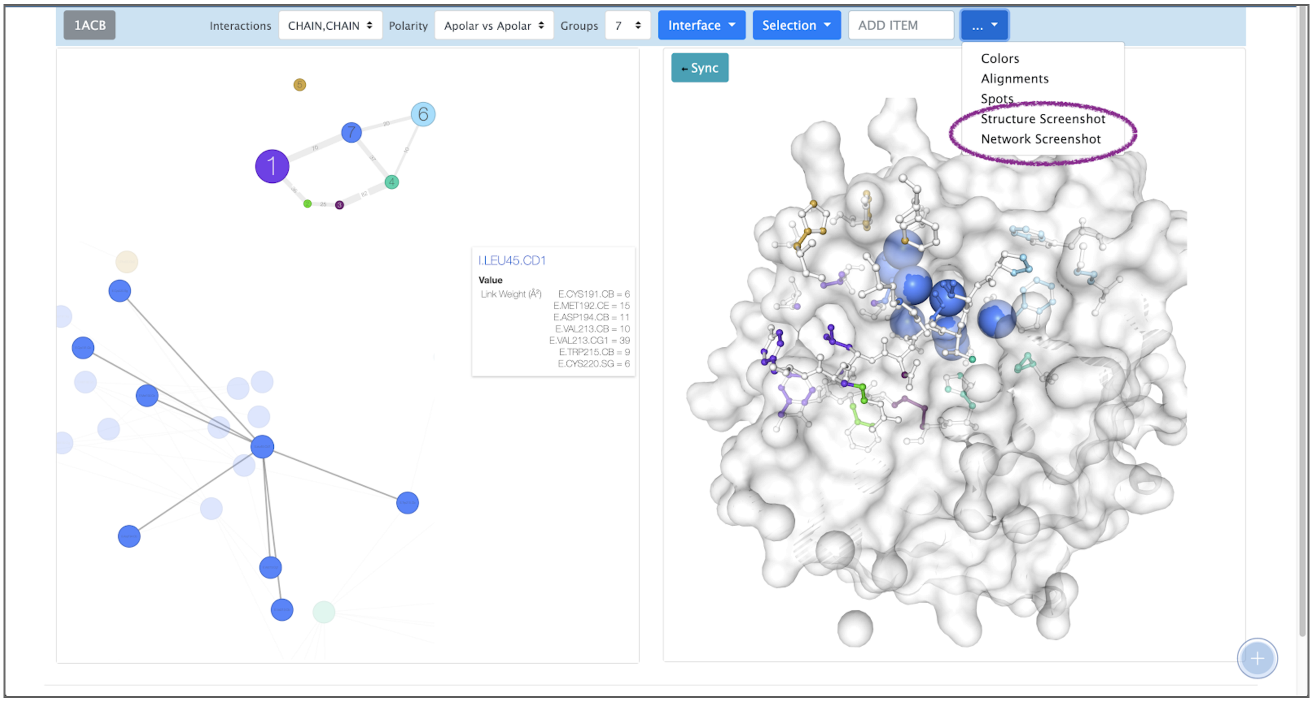


**Figure S4**. GAPIN with the possibility to take screenshots any time for both of the representations

### *How to use GAPIN*

GAPIN is a collaborative application available at https://pinga.unifei.edu.br/, which means that every time that a user imports a new PDB to GAPIN, this PDB is available for everyone. GAPIN leverages the principle of import once uses anytime.

The first step to use GAPIN is to select a PDB id it is desired to be analyzed on the top right side of the GAPIN's application as Figure S5 shows.

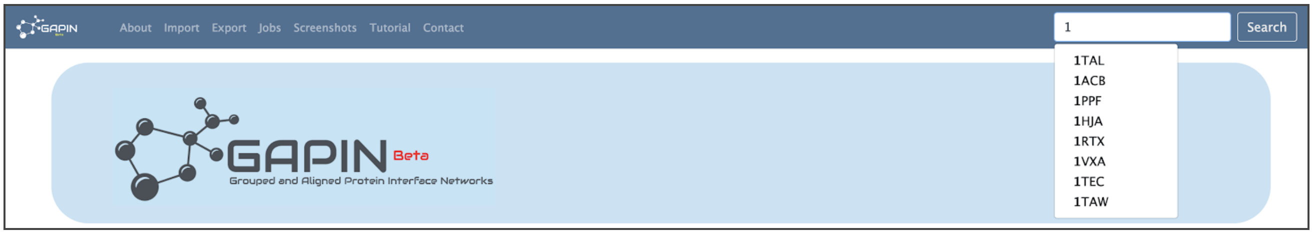


**Figure S5**. Searching for a PDB at GAPIN.

Then, two things could happen: 1) The PDB is not yet available at GAPIN, 2) The PDB selected already exists. If the first case happens, users will be redirected to the import page, Figure S6 and they will be asked to import this PDB. At this page, users can add up to five PDBs at the time separated by a comma to be imported.

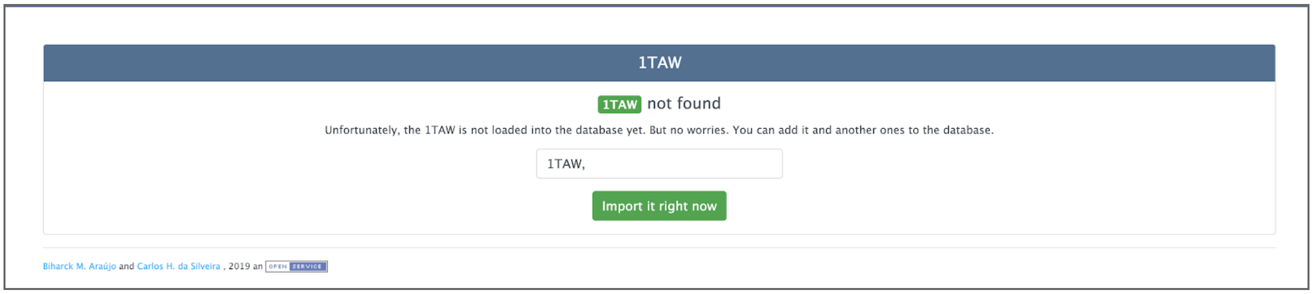


**Figure S6**: Import PDB page

The import process calls, in fact, R scripts which download PDBs from RSCB page, calculate the contact area, atom connections, group them based on the number of groups, interaction, and polarity, calculate spots and persist the output at MongoDB.

Once users start the import, the can follow the process at the Jobs page, Figure S7 and filter the table by any PDB that in under processing, Figure S8.

Show 25 entries Search:

| PDB | Date | Key | Type | Action | Percentage | Status |
|---|---|---|---|---|---|---|
| 6CYV | 2019-02-21T20:03:16.513Z | 6CYV | IMPORT | FINISHED IMPORT! | 100% | FINISHED |
| 5MLM | 2019-02-21T19:59:34.477Z | 5MLM | IMPORT | FINISHED IMPORT! | 100% | FINISHED |
| 1VXA | 2019-02-21T19:45:24.769Z | 1VXA_1B2V_LIG_AA | ALIGNMENT | FINISHED ALIGNMENT! | 100% | FINISHED |
| 1PPF | 2019-02-21T15:07:16.301Z | 1PPF_3SGB_CHAIN_AA | ALIGNMENT | FINISHED ALIGNMENT! | 100% | FINISHED |
| 1PPF | 2019-02-21T14:19:37.629Z | 1PPF_1SBN_CHAIN_AA | ALIGNMENT | FINISHED ALIGNMENT! | 100% | FINISHED |
| 1PPF | 2019-02-21T14:08:43.335Z | 1PPF_1MEE_CHAIN_AA | ALIGNMENT | FINISHED ALIGNMENT! | 100% | FINISHED |
| 1PPF | 2019-02-21T13:48:15.298Z | 1PPF_1CSE_CHAIN_AA | ALIGNMENT | FINISHED ALIGNMENT! | 100% | FINISHED |
| 1DKX | 2019-02-18T15:54:47.564Z | 1DKX | IMPORT | FINISHED IMPORT! | 100% | FINISHED |
| 2PTC | 2019-02-13T16:26:31.581Z | 2PTC_1PPF_CHAIN_ALL | ALIGNMENT | FINISHED ALIGNMENT! | 100% | FINISHED |
| 1PPF | 2019-02-11T11:52:30.907Z | 1PPF_2PTC_CHAIN_AA | ALIGNMENT | FINISHED ALIGNMENT! | 100% | FINISHED |
| 1PPF | 2019-02-11T11:50:03.200Z | 1PPF_2PTC_CHAIN_ALL | ALIGNMENT | FINISHED ALIGNMENT! | 100% | FINISHED |
| 1PPF | 2019-02-11T11:24:20.193Z | 1PPF_1FC2_CHAIN_ALL | ALIGNMENT | FINISHED ALIGNMENT! | 100% | FINISHED |
| 1CHOA | 2019-02-06T22:39:04.340Z | 1CHOA_1TEC_CHAIN_AA | ALIGNMENT | FINISHED ALIGNMENT! | 100% | FINISHED |
| 1CHOA | 2019-02-06T22:37:13.120Z | 1CHOA_1TEC_CHAIN_PP | ALIGNMENT | FINISHED ALIGNMENT! | 100% | FINISHED |
| 3SGB | 2019-02-06T14:30:43.540Z | 3SGB_1CSE_CHAIN_AA | ALIGNMENT | FINISHED ALIGNMENT! | 100% | FINISHED |
| 3SGB | 2019-02-06T14:27:18.756Z | 3SGB | IMPORT | FINISHED IMPORT! | 100% | FINISHED |
| 1CHOA | 2019-02-06T13:30:59.568Z | 1CHOA_1CSE_CHAIN_AA | ALIGNMENT | FINISHED ALIGNMENT! | 100% | FINISHED |
| 1CHOA | 2019-02-06T13:25:09.760Z | 1CHOA | IMPORT | FINISHED IMPORT! | 100% | FINISHED |
| 1SBN | 2019-02-06T12:42:26.668Z | 1SBN | IMPORT | FINISHED IMPORT! | 100% | FINISHED |
| 1CSE | 2019-02-06T12:32:39.562Z | 1CSE_1MEE_CHAIN_AA | ALIGNMENT | FINISHED ALIGNMENT! | 100% | FINISHED |
| 1MEE | 2019-02-06T12:20:55.529Z | 1MEE | IMPORT | FINISHED IMPORT! | 100% | FINISHED |

**Figure S7**. Job process logs

Show 25 entries Search: 6DGY

| PDB | Date | Key | Type | Action | Percentage | Status |
|---|---|---|---|---|---|---|
| 6DGY | 2019-01-14T20:15:13.871Z | 6DGY | IMPORT | FINISHED IMPORT! | 100% | FINISHED |

Showing 1 to 1 of 1 entries (filtered from 72 total entries) Previous 1 Next

**Figure S8**. Jobs process logs filtered by a PDB

When the process finishes, the status will change to finished and users can start working with the desired PDB.

In case of the PDB already exists or has been imported, users are redirected to the main page, Figure S9, where they can start interacting with the PDB. At the right side, Figure S9-C, users can interact with the interface structure built with NGL such as rotate to any side with the mouse left click, either zoom-in or zoom-out holding the mouse right click and scrolling up or down, reposition the structure and any part of the layer with the mouse middle click.

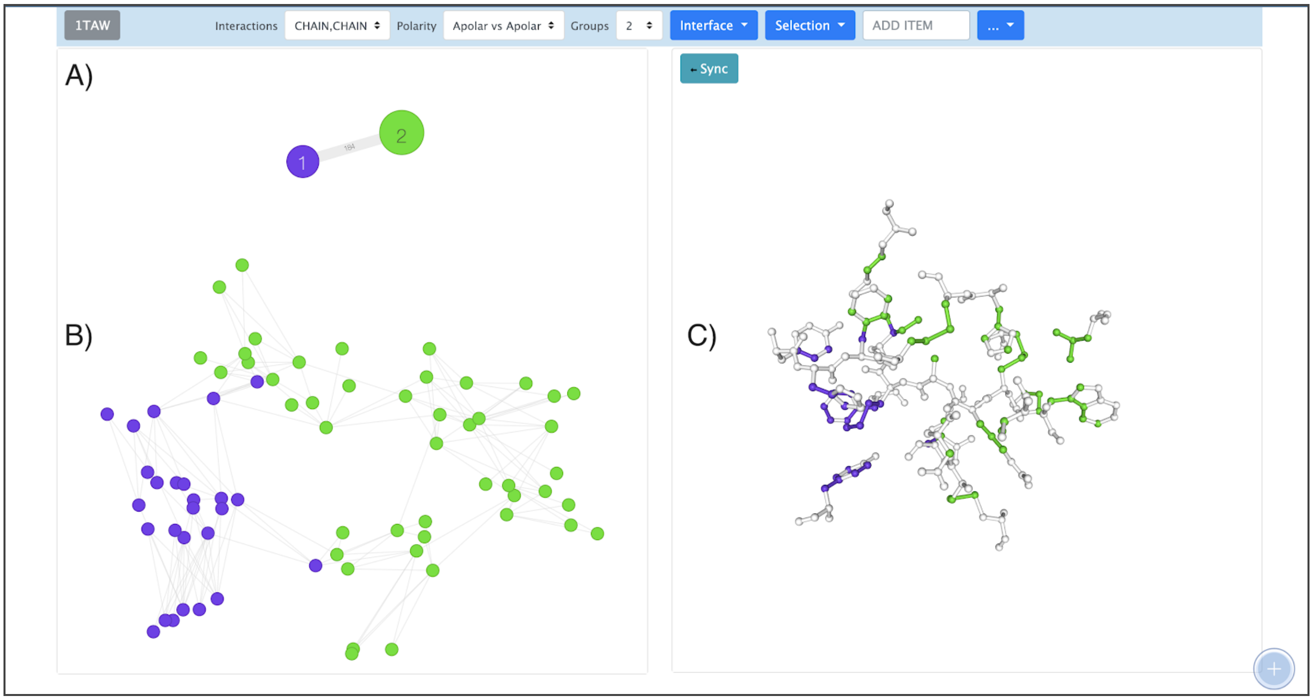


**Figure S9**. GAPIN's main interaction page

At the left down side, Figure S9-C, users can interact with the atom connections network built with D3Plus where each node represents an atom and the edges represent the contact area among the atoms. at the left top side, Figure S9-A, a called super-cluster graph shows the high-level representation of the connections network graph right below if it. The node round size means the node weight, going deeper, means the sum of all contacts are from the nodes that belong to its group and the edges the sum of the connections.

Users can select any atom or group lift clicking the networks graphs to see what that connection is on the structure side, Figure S10 and S11.

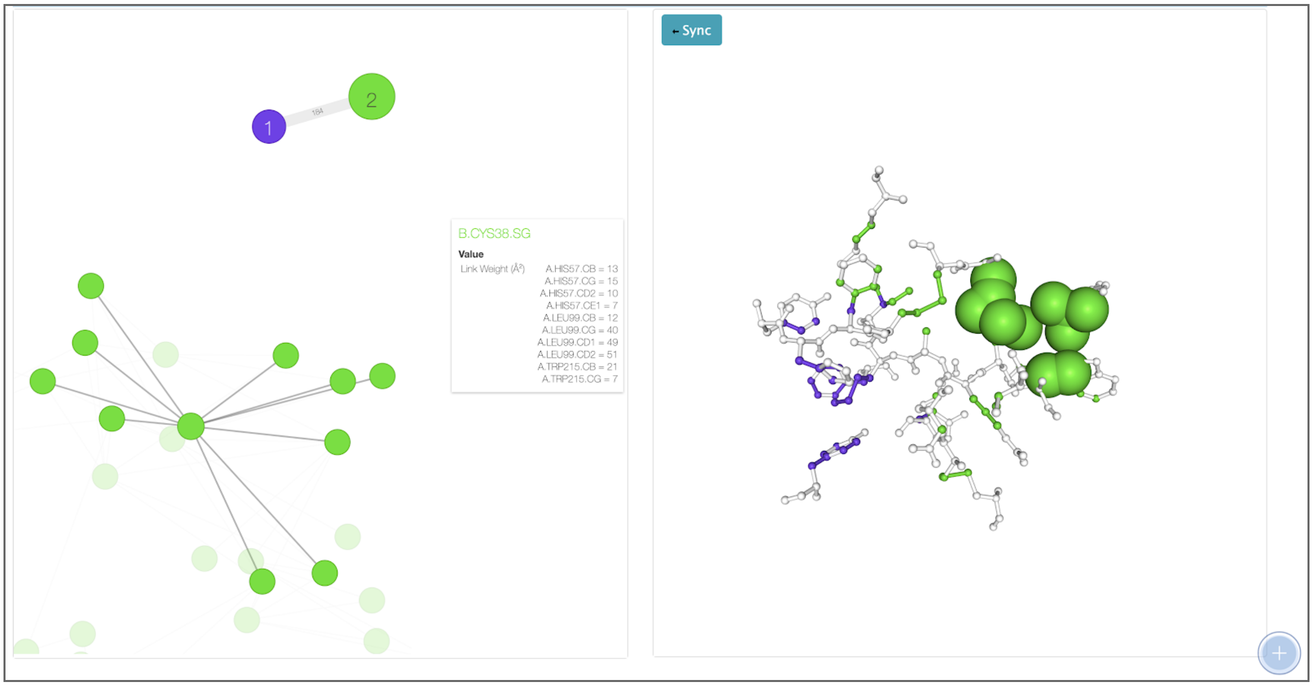


**Figure S10**. Node selection at the atom network graph

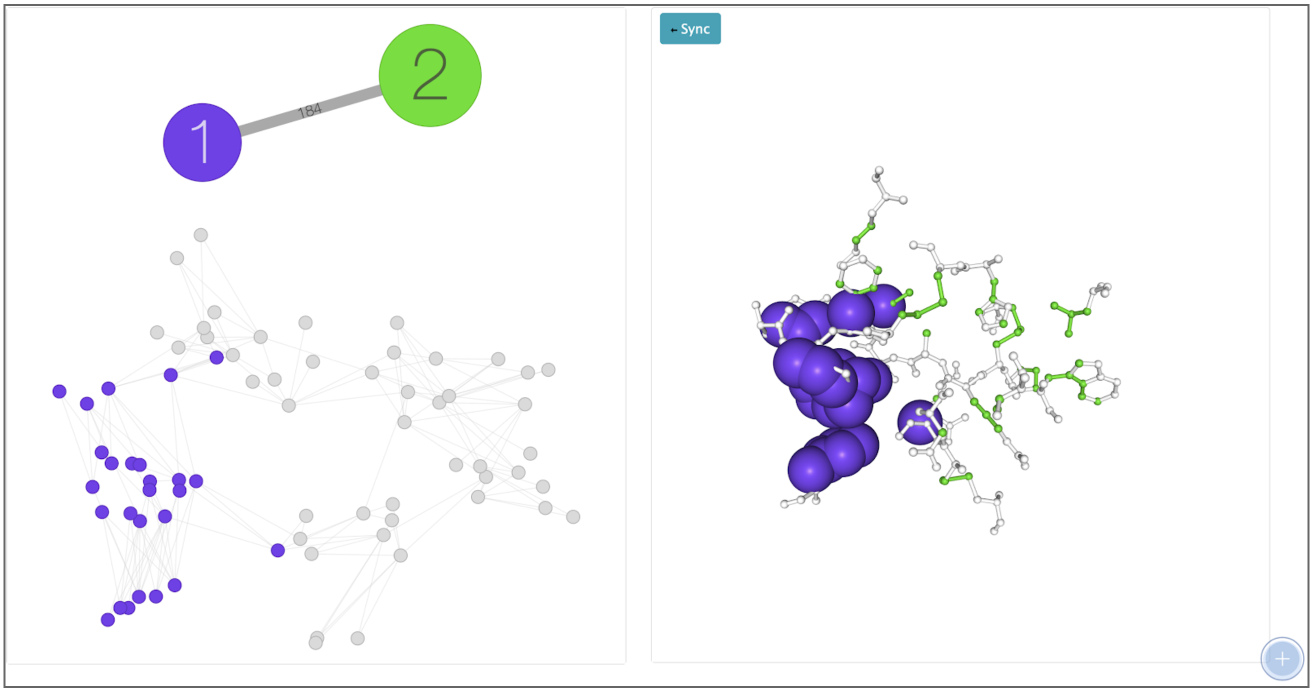


**Figure S11**. Group 1 selected at the super-cluster graph being reflected at the structure on the right side and highlighted at the atom network graph.

At the top bar, Figure S12, right above of the network and structure representations, there is a set of options users can use in order to analyze better the macromolecule.

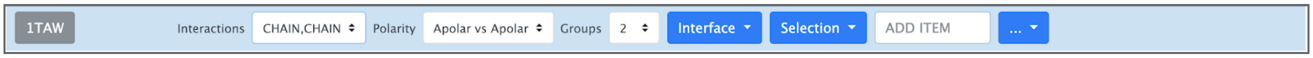


**Figure S12**. The toolbox that is applied to the interface structure and contact graph

In interactions, users can choose between CHAIN vs CHAIN interactions, which means all contacts found at the surface level of the macromolecule interacting to each other from their chains, or ANY, LIG which means the interactions between atoms from the macromolecule interacting with Ligands. The second option is the Polarity. Polarity means that users can analyze the macromolecule considering only Apolar contacts, Polar contacts or them all. The third option is Groups. At the first time users load the PDB, the default configuration is to show the networks and structure divide by two groups, but users can select other combinations up to ten[1]. The algorithm that calculates the groups is based on Spectral Clustering (Luxburg, U., 2007).

Right after the Groups, there is a set of tools that are applied to the interface structure such as Structure representation, the color format, Visualization and the ability to add the macromolecule surface. Figure S13 shows a combination of this set of tools applied to the PDB 1PPF (Chakrabarty, B., & Parekh, N., 2016).

[1] Due to the size of some macromolecules they may not reach 10 groups.

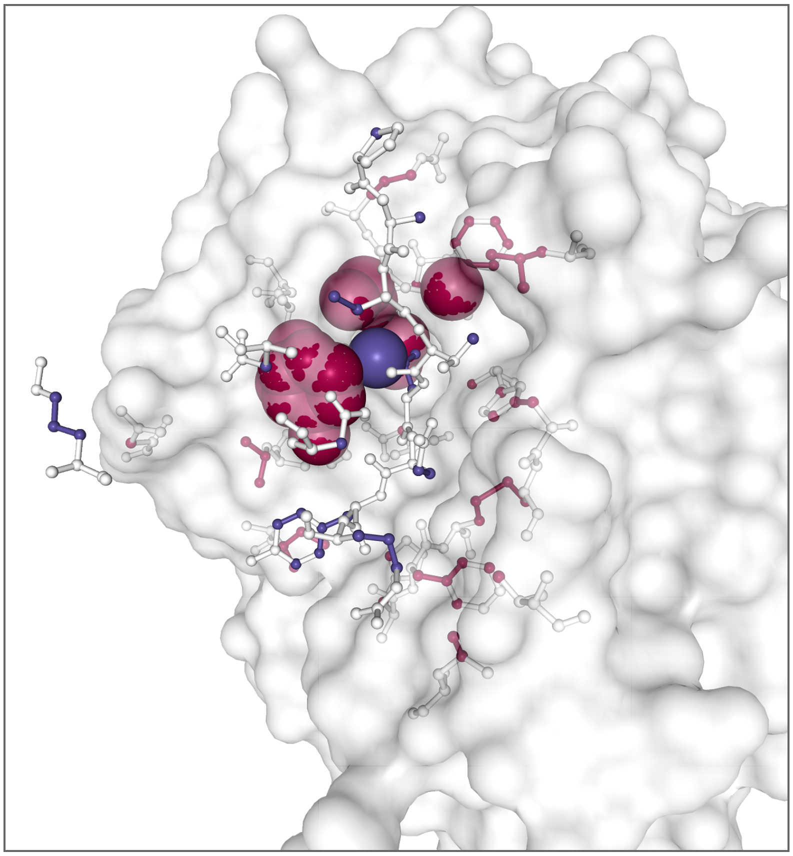

**Figure S13**. Using the interface toolbox to get a better visualization for the analyzed structure

Right after interactions toolbox, there is the selection toolbox which basically does almost the same thing as interactions toolbox but for the selection. So, every time users select an interaction at the network graph level, users can use this pallet to change the representation to either ball stick or spacefill, visualize the full residues in white, change the colors of the selection based on the number of groups, polarity, elements or interactions and finally clear the selection changes.

On the right side of selection toolbox, there is an input field where users can search for an specifically desired atom and see it at the structure right bellow of it.

The last menu item with the three dots adds options like the ability to change the color of the groups, Figure S14.

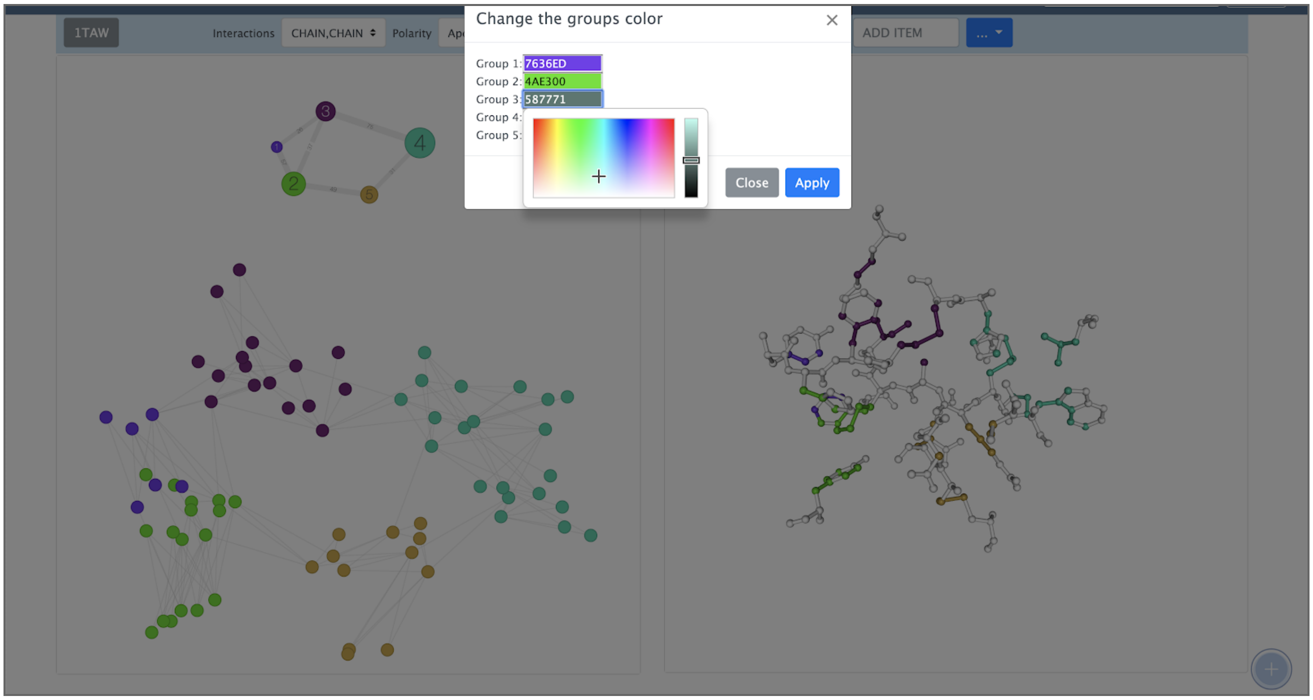


**Figure S14.** The ability to change the colors of the groups any time

There is also the spots representation for the selected macromolecule, Figure S15, where users can see how strong is the connection throughout the number of groups increases

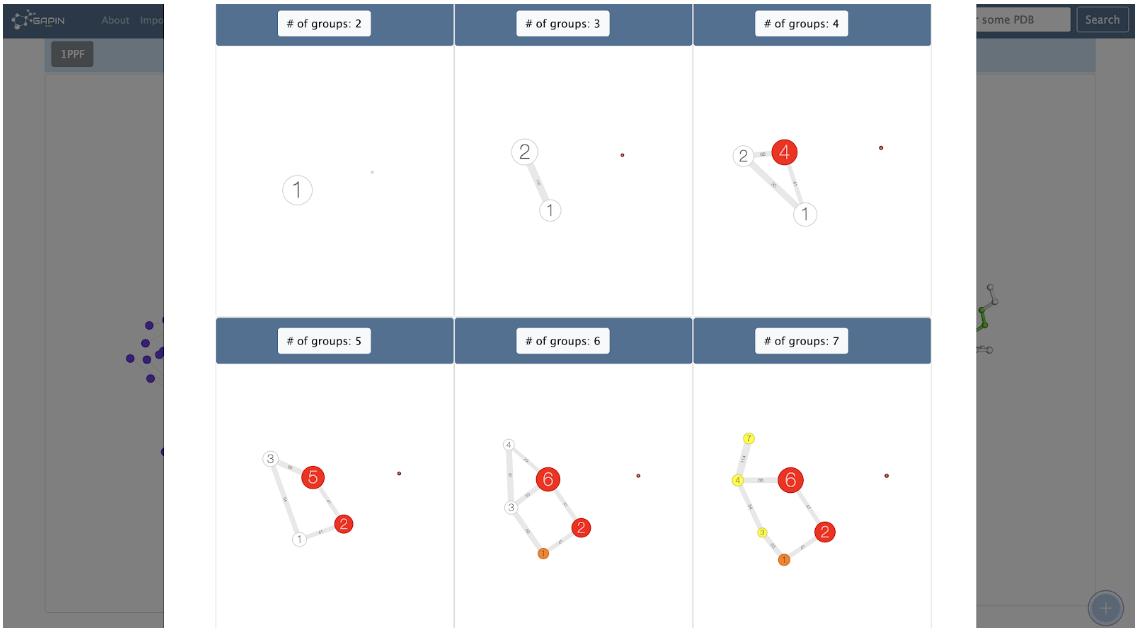


**Figure S15**. Spots representation for the selected protein

At the spot representation users can see that as more red or hot is the group, stronger is to break it alongside increases the number of groups. And the opposite is, as more white or cold is the group, it is easier to break this cluster.

There is also the possibility to take and download unlimited snapshots for all representations any time as mentioned before.

Finally, at this menu, there is an option called alignments. The alignment section allows users to automatically align any macromolecule to any other macromolecule available at the GAPIN's database.

The same idea of import is applied here where if there is no alignment yet for the users choice, they can start the alignment process, Figure S16, and follow the Job at the Job section, Figure S17.

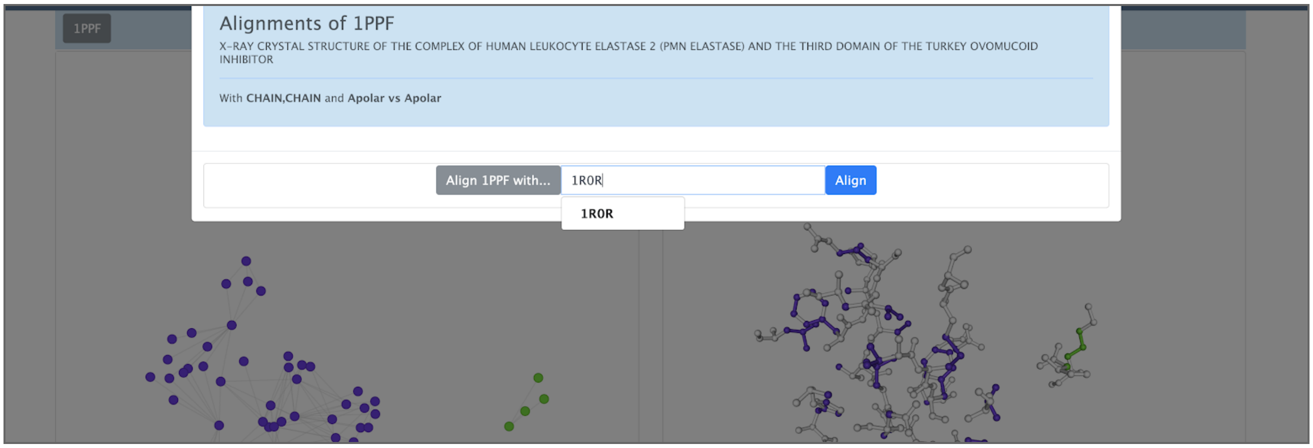


**Figure S16**. Starting the alignment between two PDBs

Show 25 entries                Search: 1ppf

| PDB | Date | Key | Type | Action | Percentage | Status |
|---|---|---|---|---|---|---|
| 1PPF | 2019-02-21T22:36:12.034Z | 1PPF_1A22_CHAIN_AA | ALIGNMENT | Alignment has been started | 0% | |
| 1PPF | 2019-02-21T15:07:16.301Z | 1PPF_3SGB_CHAIN_AA | ALIGNMENT | FINISHED ALIGNMENT! | 100% | FINISHED |
| 1PPF | 2019-02-21T14:19:37.629Z | 1PPF_1SBN_CHAIN_AA | ALIGNMENT | FINISHED ALIGNMENT! | 100% | FINISHED |
| 1PPF | 2019-02-21T14:08:43.335Z | 1PPF_1MEE_CHAIN_AA | ALIGNMENT | FINISHED ALIGNMENT! | 100% | FINISHED |
| 1PPF | 2019-02-21T13:48:15.298Z | 1PPF_1CSE_CHAIN_AA | ALIGNMENT | FINISHED ALIGNMENT! | 100% | FINISHED |
| 2PTC | 2019-02-13T16:26:31.581Z | 2PTC_1PPF_CHAIN_ALL | ALIGNMENT | FINISHED ALIGNMENT! | 100% | FINISHED |
| 1PPF | 2019-02-11T11:52:30.907Z | 1PPF_2PTC_CHAIN_AA | ALIGNMENT | FINISHED ALIGNMENT! | 100% | FINISHED |
| 1PPF | 2019-02-11T11:50:03.200Z | 1PPF_2PTC_CHAIN_ALL | ALIGNMENT | FINISHED ALIGNMENT! | 100% | FINISHED |

**Figure S17**. Jobs page to follow the alignment process

At the end of the alignment process, users can see the alignment such as Figure S18.

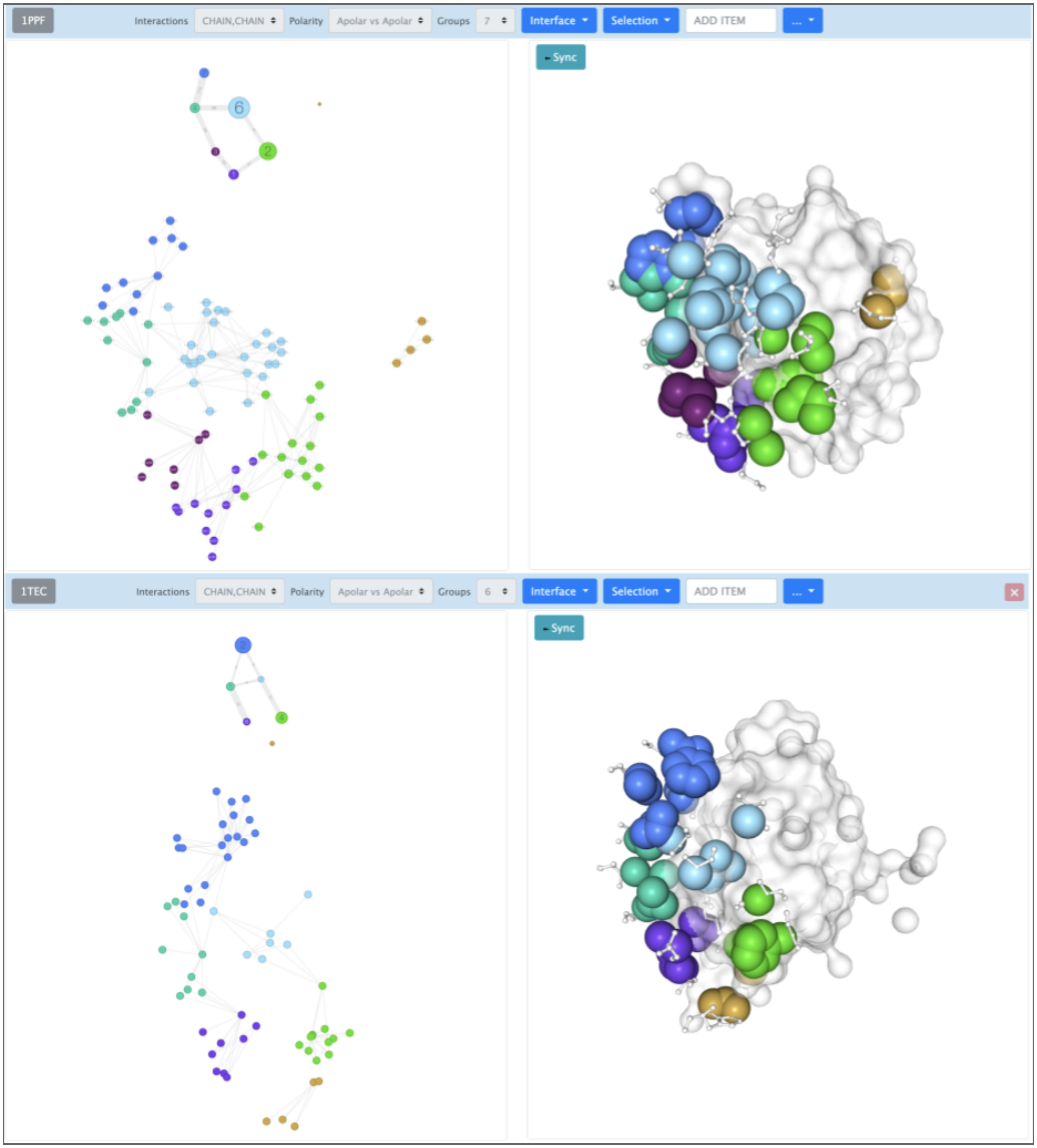


**Figure S18**. 1PPF and 1TEC aligned

Users are allowed to change the number of groups to get the best combination they desire. One import issue is, every time the number of groups change, the way that GAPIN presents the network structure is based on the original structure. So, if the users want to see the network aligned, they just have to click at the sync button and the network will follow the structure position.

Furthermore, GAPIN also allows users to align any structure manually and synchronize the changes in the structure with the graph at the left side with just one click, Figure S19 shows the structure rotate but the network not synchronized and Figure S20 shows after synchronization.

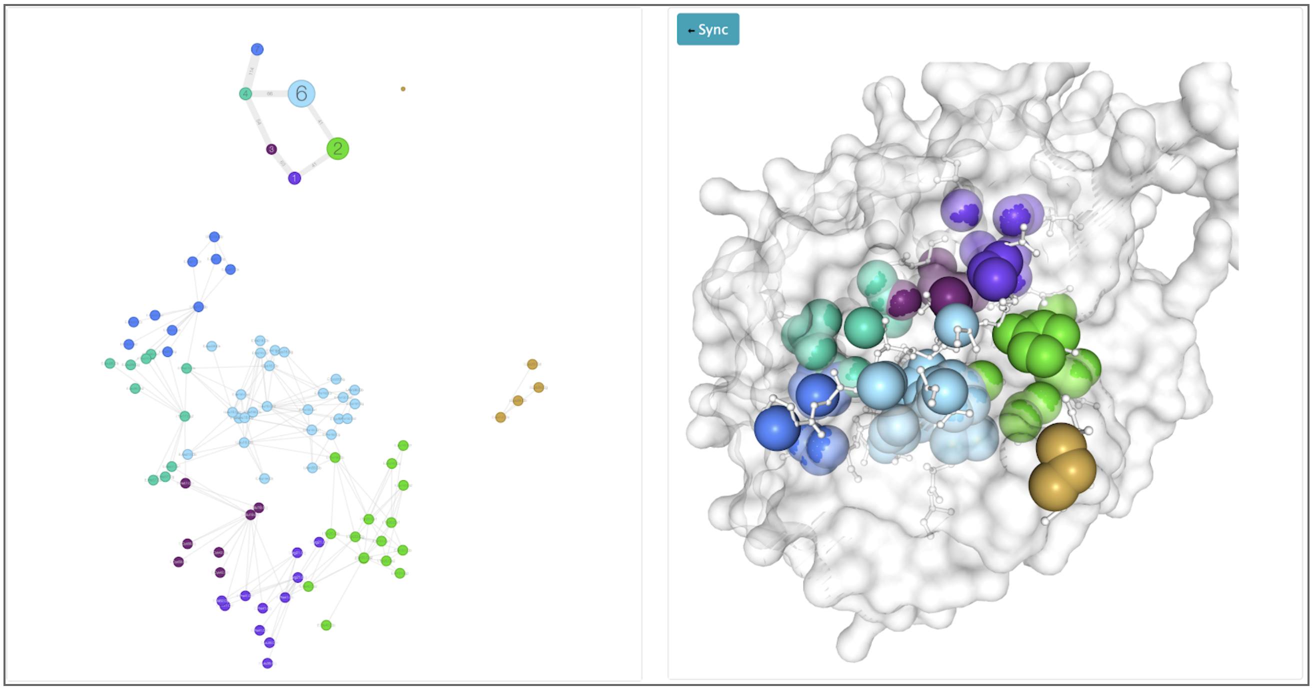


**Figure S19**. Structure rotated and out of sync with the network graph

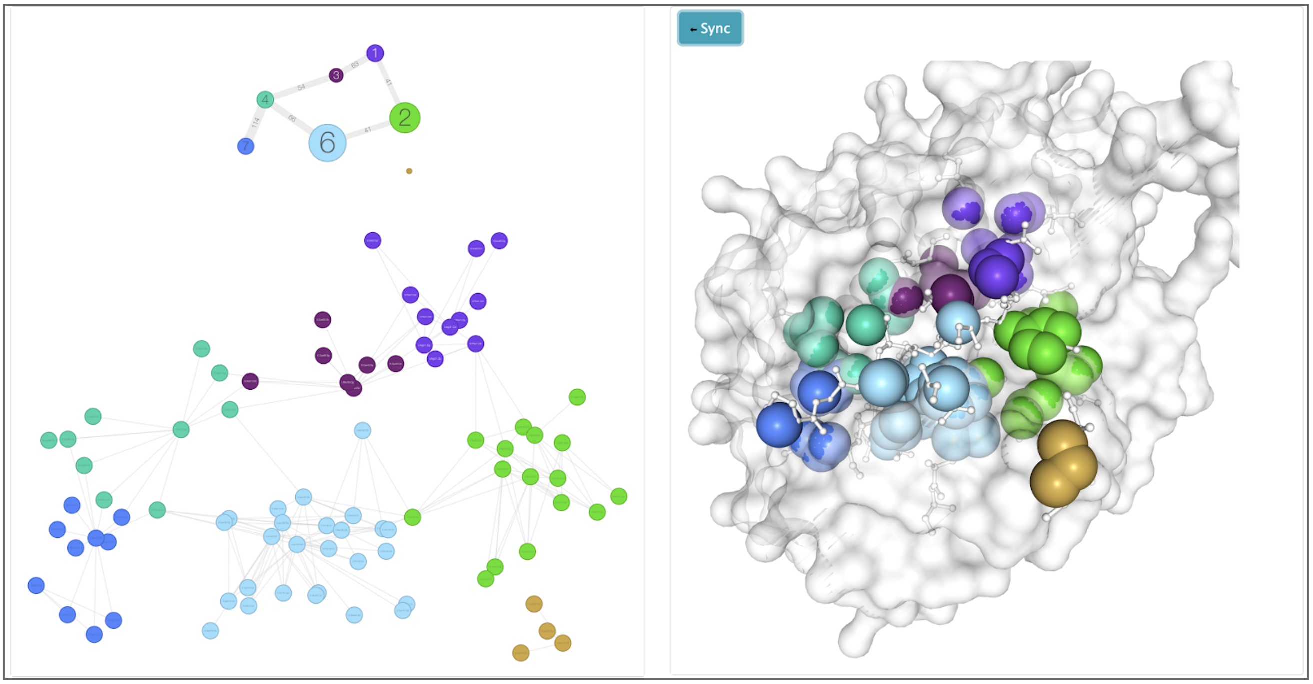


**Figure 20**. Structure rotated and synchronized with the network graph

GAPIN also allows users to add as many macromolecules they desire to analyze using the plus icon, Figure S21, at the bottom right side and the macromolecules will be distributed one bellow to another.

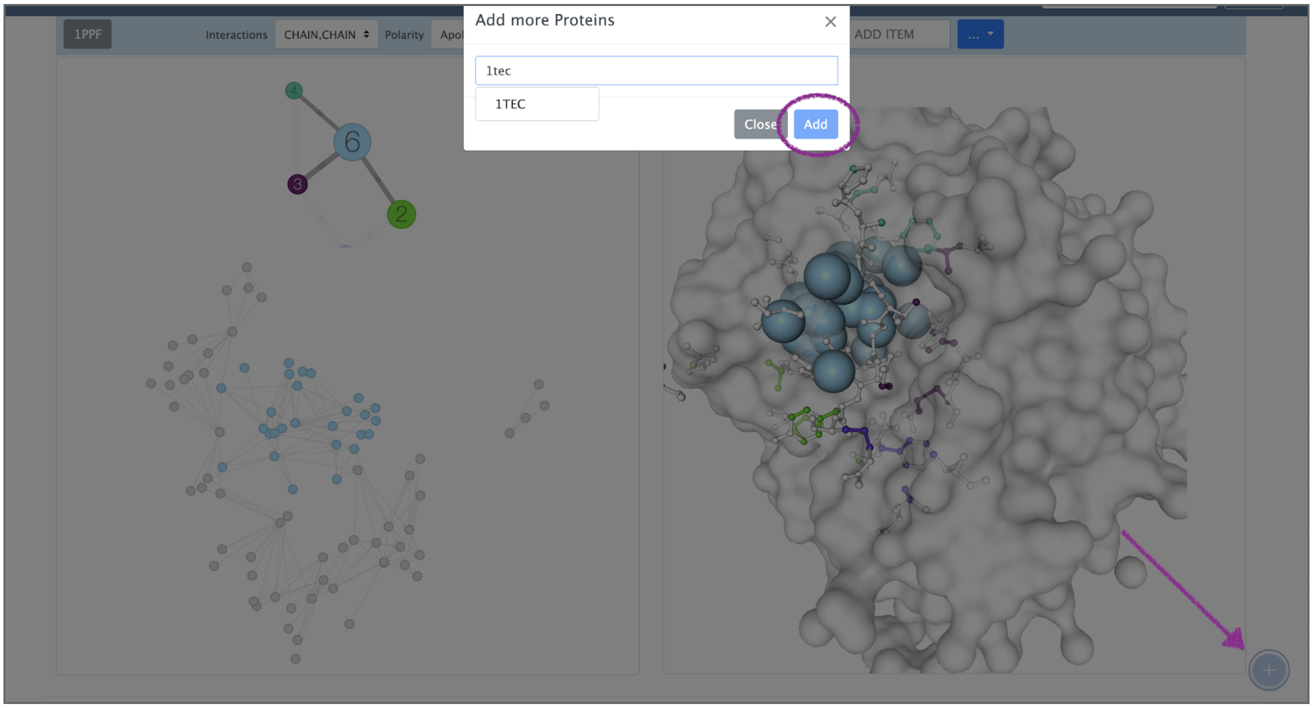


**Figure S21**. GAPIN allows users to add as many macromolecules they want to

### *References*


Duvall, Paul M., Steve Matyas, and Andrew Glover. Continuous integration: improving software quality and reducing risk. Upper Saddle River, NJ: Addison-Wesley, 2007. Print.

Humble, J., 2010. Continuous Delivery: Reliable Software Releases through Build, Test, and Deployment Automation (Addison-Wesley Signature Series (Fowler)). Addison-Wesley.

Bitbucket. (2012). Retrieved January 12, 2019, from https://bitbucket.org/

Bitbucket Pipelines. Retrieved January 12, 2019, from https://bitbucket.org/product/features/pipelines

Casciaro, Mario. Node.js design patterns: get the best out of Node.js by mastering a series of patterns and techniques to create modular, scalable, and efficient applications. Birmingham: Packt Publishing, 2016. Print.

Gallaba, K., Mesbah, A., Beschastnikh, I. (2015). Don't Call Us, We'll Call You: Characterizing Callbacks in Javascript. 2015 ACM/IEEE International Symposium on Empirical Software Engineering and Measurement (ESEM) : proceedings::-23 October 2015, Beijing, China. Piscataway, NJ: IEEE, 2015. Print.

NodeJS. Retrieved January 13, 2019, from https://nodejs.org/en

ExpressJS. Retrieved January 13, 2019, from https://expressjs.com

R Core Team (2018). R: A language and environment for statistical computing. R Foundation for Statistical Computing, Vienna, Austria. URL https://www.R-project.org/.

MongoDB. Retrieved January 13, 2019, from https://www.mongodb.com

Otto, M., & Thornton, J. (2015). Bootstrap· The world's most popular mobile-first and responsive front-end framework.'. *Getbootstrap.com*.

HandleBars JS. Retrieved January 13, 2019, from https://handlebarsjs.com

Rose, A. S., Bradley, A. R., Valasatava, Y., Duarte, J. M., Prlic, A., & Rose, P. W. (2018). NGL Viewer: Web-based molecular graphics for large complexes. *Bioinformatics*, *1*, 4.

D3plus. (n.d.). Retrieved December 30, 2018, from https://d3plus.org/

DataTables. (2008). Retrieved December 30, 2018, from https://www.datatables.net/

Kouranov, A., Xie, L., de la Cruz, J., Chen, L., Westbrook, J., Bourne, P. E., & Berman, H. M. (2006). The RCSB PDB information portal for structural genomics. *Nucleic acids research*, *34*(suppl_1), D302-D305.

Luxburg, U. (2007). A tutorial on spectral clustering. Statistics and Computing, 17(4), 395–416.

Chakrabarty, B., & Parekh, N. (2016). NAPS: Network analysis of protein structures. Nucleic Acids Research, 44(W1)
