## Supplementary material for "GAPIN: Grouped and Aligned Protein Interface Networks"

[1] Bioinformatics PhD Program, Universidade Federal de Minas Gerais, 31270-901, Brazil.

[2] School of Medicine - EMED / Universidade Federal de Ouro Preto, 35400-000, Brazil.

[3] Department of Computer Science, Universidade Federal de Viçosa, 36570-900, Brazil.

[4] European Molecular Biology Laboratory, European Bioinformatics Institute, Wellcome Genome Campus, Hinxton, UK.

[5] Institute of Applied and Pure Sciences, Universidade Federal de Itajubá, 35903-087, Brazil.

[6] Department of Computer Science, Universidade Federal de Minas Gerais, 31270-901, Brazil.

[7] Institute of Technological Sciences, Universidade Federal de Itajubá, 35903-087, Brazil.

#### *1) Supplementary Material and Methods*

##### Graphs

**Lower-level graphs:** adjacency matrices are built having atoms as nodes and contact areas as weight for the edges. Only contacts between atoms of different chains (interchain contacts) are considered, producing bipartite undirected graphs. Ligands, including waters and ions, are considered as chains. Hence, in algorithms terms, a chain is a label that groups related atoms.

**Higher-level graphs**: lower-level graphs are clustered into higher-level graphs in order to introduce network modularity analysis. In a modular graph context, each node is a cluster whose vertex label is the sum of contact areas among atoms of the same group, and the edge label is the sum of contact areas among atoms of different groups.

Examples of lower-level and higher-level graphs can be seen in figure S2.

##### Node centroids and visual synchronization

A centroid with 3D coordinates is associated to each node in both the lower-level and higher-level graphs. For lower-level nodes, the centroids are the 3D coordinates of corresponding atoms. For higher-level nodes, the centroids are the geometric centers of clustered atoms. For both graph views, it is applied a layout that takes into account these coordinates, projected on the x-y plane. This allows synergistic manipulations and visual synchronizations between graphs and rendered PDB structures. It is important to add that for both views, nodes and edges are drawn proportionally with their respective contact areas. See Tutorial document for usage details.

#### Community detection algorithm

It is used a normalized spectral clustering algorithm rather similar to those described in (Luxburg, 2007) but with an asymmetric Laplacian matrix ($L_{rw}$) built from the adjacency matrices of lower-level graphs, and partition around medoids (PAM) as final clustering algorithm applied over the eigenvector basis. A graph can be divided into disjoint partitions (clusters) by cutting the appropriate edges. Generally, good clusterings are those that aggregate nodes and edges in groups with minimal cuts. By Rayleigh-Ritz theorem (Luxburg, 2007), with $L_{rw}$ we can generate an approximation of a special minimal cut called Ncut, in which cuts are normalized by the sum of partition edge weights. So, minimize Ncut is equivalent to make graph partitions (clusters) with maximum edge weights inside the groups and minimum outside (For mathematical details, see Luxburg, 2007).

In addition to its simplicity, good performance and robust benchmark scores on small networks (Lancichinetti and Fortunato, 2009; Newman, 2004), another important factor in choosing spectral clustering as community detection algorithm is the possibility of easily implementing a cluster scanning (see the cluster scanning section below).

#### Clustering quality

The quality of spectral clustering is evaluated following the intuition of Ncut minimization described above: a better clustering is one that maximizes contacts internally to the group and minimizes externally. Looking at a higher graph adjacency matrix (like in Example 1S), it is easy to see that good clusters are those that concentrate the values on the diagonals.

So, a simple quality metric is to take the ratio between the node inner area over the node total area. In Example 1S, the cluster quality for node 1 would be: 335.91/(29.14+80.99+335.91) = 0.75. That is, 75% of the node total area involve edge weights between atoms belonging to it.

It is noteworthy that a perfect clustering with maximum quality would produce a diagonal adjacency matrix `n` x `n`, with `n` connected components. Interestingly, this higher graph adjacency matrix can be seen as a confusion matrix (Zaki and Meira, 2016), used to measure the performance of an algorithm in some machine learning techniques. A perfect classifier would also generate a diagonal contingency table with all true-positive (or true-negative) values in that diagonal (and null for all remaining values). In this sense, most of the metrics derived from the confusion matrix may be valid for higher graph adjacency matrix, such as precision, recall, accuracy, F1 score, etc, with the difference that the latter is symmetric. This causes false-negative values to be equal to false-positive, unifying several of confusion matrix metrics, such as recall and precision. Our quality metric, for example, can be calculated as a sensitivity (recall) score.

**Example S1**: higher graph adjacency matrix

|  | [,1] | [,2] | [,3] | [,4] | [,5] |
|---|---|---|---|---|---|
| [1,] | 335.91 | 80.99 | 29.14 | 0.00 | 0.00 |
| [2,] | 80.99 | 470.10 | 55.42 | 0.00 | 62.60 |
| [3,] | 29.14 | 55.42 | 1545.08 | 43.52 | 0.00 |
| [4,] | 0.00 | 0.00 | 43.52 | 986.60 | 41.30 |
| [5,] | 0.00 | 62.60 | 0.00 | 41.30 | 310.19 |

#### Cluster scanning

In partition clustering algorithms, as the number of clusters (`k`) must be specified in advance, the usual approach is looking for a `k` that maximizes the overall cluster quality, such as silhouette coefficient (Rousseeuw, 1987) or modularity measure (Newman, M. E. J.; Girvan, M., 2004). In this work, we follow a scanning method (da Silveira et. al., 2009) showing the graph partitions in successive `k`. This may bring some valuable information about the structural organization of the graph, since the clustering algorithm is trying to minimize the Ncuts for each `k`. It also allows the user to know what the propensity of the node under analysis to be partitioned in the next `k`. In Figure S1, it is possible to see a sequence of k-partitioning (from 3 to 6) for the nonpolar-nonpolar higher graph of 1PPF. Nodes in red were more preserved from partitioning than others. The increment of `k` stops when the minimum clustering quality reaches 50% or k reaches 20 partitions (this limit of 20 was imposed due to performance issues).

#### Preservation index

We create a preservation index that measures how much a node resists to be repartitioned. Given a sequence of successive k-partitions, for each corresponding node between partitions i and i+1, the index returns the average area ratio ($A_{i+1}/A_i$) if this ratio is greater than a given threshold (defined as 0.85). Corresponding nodes are those whose centroids are less than 1.4 angstroms apart. If the ratio is less than the threshold or the distance is greater than 1.4, then the ratio is reset to zero (see equation 1). Example 2S show how this index works with 2 interactions.

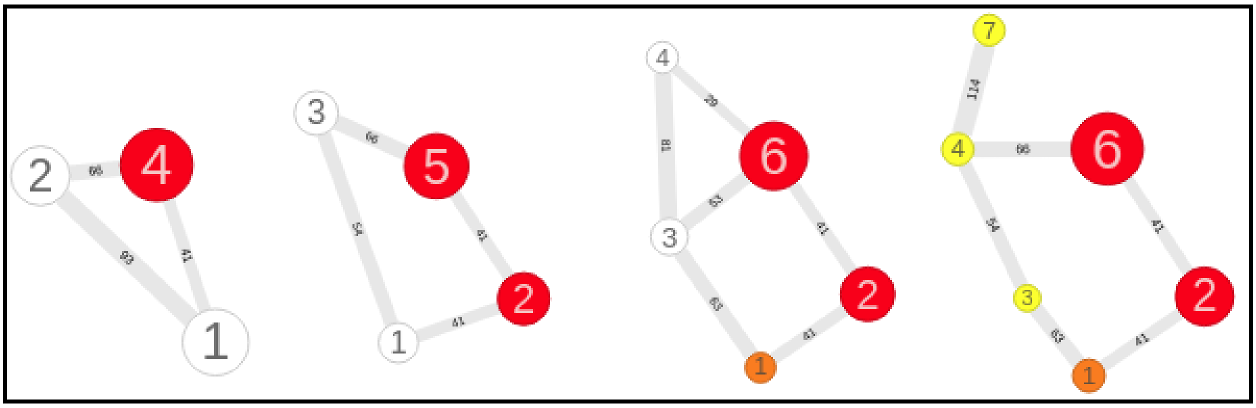

**Fig. S1.** Cluster scanning for nonpolar-nonpolar higher graph in 1PPF. Colors indicate the preservation index, going from low (white-yellow) to high (red-brow).

**Example S2**: preservation index building

[[1]]

| | V3 | V4 | Dist | A3 | A4 | A4/A3 | Preserve_idex |
|---|---|---|---|---|---|---|---|
| 1-1 | 1 | 1 | 0.002 | 1242.17 | 1242.17 | 1.00 | 1.00 |
| 2-2 | 2 | 2 | 2.197 | 2724.62 | 2095.24 | 0.00 | 0.00 |
| 3-3 | 3 | 3 | 0.573 | 571.05 | 571.05 | 1.00 | 1.00 |

[[2]]

| | V4 | V5 | Dist | A4 | A5 | A5/A4 | Preserve_idex |
|---|---|---|---|---|---|---|---|
| 1-1 | 1 | 1 | 0.002 | 1242.17 | 1242.17 | 1.00 | 1.00 |
| 2-2 | 2 | 2 | 0.718 | 2095.24 | 2026.27 | 0.97 | 0.48 |
| 3-3 | 3 | 3 | 0.002 | 571.05 | 571.05 | 1.00 | 1.00 |
| 4-4 | 4 | 4 | 0.002 | 450.20 | 450.20 | 1.00 | 0.50 |

Where:
$V_i$ = vertex with index i
Dist = distance between $V_i$ and $V_{i+1}$
$A_i$ = area of node i
$A_{i+1}/A_i$ = area ratio
Preserve_id = preservation index

#### Contact area

It is computed for each atom pairs, using Biharck-Alves-Romanelli-Silveira (BARS) heuristic. This contact area is equivalent to the excluded or untouched area from a rolling water molecule (probe) between two isolated atoms of different chains. So, a null contact area indicates that one (or more) water molecule could interpose between two atoms, defining a potential cavity in terms of the heuristic. As a welcome consequence, atoms of different chains with non-zero contact areas delimit the interface between biomolecules.

One of advantages of BARS methodology is the fact that the contact areas can be computed analytically by an equation (2) instead of a procedure as done in nAccess (Lee and Richards, 1971; Hubbard and Thornton, 1993). This saves considerable computing time, since the contact area between two atoms can be computed in order of one - $O(1)$. Despite this simplification, BARS calculated areas still retain significant correlations with solvent-accessible surface areas (ASAs). This can be seen in figure S5, built with a sample of 68 chain-chain complexes from Affinity Database 2.0 (Kastritis et. al., 2011).

The BARS equation can be described as:

$$A(R_1, R_2, d) = 2\pi(R_1^2 + R_2^2) - \pi(R_1 + R_2)d\left[1 + \left(\frac{R_1 - R_2}{d}\right)^2\right] \qquad (2)$$

where: $A$ is the contact area, $R_1 = r_1 + p$ and $R_2 = r_2 + p$, with $r_1$ and $r_2$ as the van der Waals radii from atoms $a_1$ and $a_2$ ,respectively; $p$ is the radius of probe (set default to 1.4); $d$ is the Euclidean distance between atoms $a_1$ and $a_2$, assuming $r_1 \geq r_2$. The demonstration of this equation can be found in Appendix A, at the end of this document.

#### Atom interactions

In the current version, GAPIN works with two types of atomic interaction: polar and nonpolar. Atoms are classified according to Table S01, which was adapted from (Sobolev et. al., 1999). It was chosen to be more conservative regarding hydrophobic interactions at protein-protein interfaces (PPI), classifying some protein carbons as polar. In this way, all backbone atoms are set as polar.

**Table S1.** Binary atomic classification according to polarity.

| ATOM | POLARITY |
|---|---|
| Protein backbone carbons (CA and C) | Polar |
| Protein carbons in a covalent bond to polar atoms (except for HIS and TRP)* | Polar |
| Other protein carbons | Nonpolar |
| Protein CYS.S in SS-bond, MET.S | Nonpolar |
| Any O or N | Polar |
| Any non-protein carbons | Nonpolar |
| Any other atoms | Polar |

* given its intrinsic behavior in hydrophobic cores (White and Wimley, 1999).

#### Higher graph alignments

Alignments of higher graph can be done manually or automatically. For the first case, the user loads two (or more) PDBs and operates the alignment with the mouse, using the synchronize button between rendered PDB structures and graphs. For the second case, after load an initial PDB, the user must select the alignment option from the menu, and choose another PDB to align with. Whenever possible, two list of graphs will be returned with two levels of graph clustering quality and best scores associated to them. For more usage details, see the Tutorial document. See also section 3 below on how the figures in this publication were generated, specially figure 1A.

**Algorithm 1.** Algorithm `Topos` for graph alignments

**Input:**

```
adjacency matrices A1, A2; centroid matrices C1, C2
```

**Output:**

```
a list of the best transformation matrices that aligns C2 into C1
```

```
Function ALIGNMENT(A1,A2,C1,C2)
  n1 = number of nodes in C1
  n2 = number of nodes in C2
  k = number of nodes in a subgraph
  n = number of right singular vectors
  m = size of best list
  ListC1 = node combinations ids(n1,k)
  ListC2 = node combinations ids(n2,k)
  Best = empty list of size m
  For each i in ListC1 do
    V1 = n first right singular vectors of SVD(C1,ListC1[i])
    For each j in ListC2 do
      V2 = n first right singular vectors of SVD(C2,ListC2[j])
      Rot = align vectors by direction cosines (V1,V2)
      Score = EVALUATE(A1,A2,C1,tranformation(C2,Rot))
      Best = save best scores(Best,Score,Rot)
  Return(Best)
```

```
Function EVALUATE(A1,A2,C1,C2)
  ListD = list of the corresponding atoms (C1,C2)
  ListA = list of the area ratio of superimposed nodes (A1,A2,ListD)
  ListE = list of the cossine of superimposed edges (C1,C2,ListD)
  return(sum(ListA)+sum(ListE))
```

Our automatic alignment algorithm (called `Topos`) takes into account both the graph topology and positions of node centroids, such as described by the pseudocode (Algorithm 1). The pivotal idea behind this algorithm is the alignment of `n` first right singular vectors, that results from the singular value decomposition (SVD) of centroid matrices. SVD is a generalized factorization for any matrix $\mathbf{A}$, given by $\mathbf{A} = \mathbf{LSR}^t$, where $\mathbf{L}$ is called the left singular vector matrix, $\mathbf{S}$ is a diagonal matrix with singular values, and the columns of $\mathbf{R}$ matrix (rows of $\mathbf{R}^t$) have the right singular vectors (Zaki and Meira, 2016). As $\mathbf{L}$ and $\mathbf{R}$ are orthogonals, it is possible to use both as basis vectors. The alignment of two different basis vectors can be made by direction cosines of the angles between each corresponding axis (Kelly, 2019).

EVALUATE is a function that scores the alignment taking into account the ratio of the areas between the superimposed nodes and the cosine of the angles between the overlapping edges. The higher the score, the better.

It is possible to search subgraphs that better align two graphs adjusting the parameter k (number of nodes in a subgraph), but this generates a combinatorial process. As mitigating factors, in addition to the fact that higher graphs have a few nodes, there is a limit for the maximum number of partitions (currently, 20). Thus, this combinatorial process can be computationally tractable.

A detailed analysis of this algorithm is beyond the scope of this paper. Another publication is being prepared specifically for this purpose. Anyway, the automatic alignment is not essential for the results presented here, since it could also be done manually.

In the current version of GAPIN, only pairwise interface alignments (two PDBs at a time) are allowed. It does not do mass alignments of PDBs yet. Chain-ligand interface alignments are not enabled either, but they will be available soon.

#### 2) Supplementary Experiment Data and Results

**Table S2.** Set of serine-peptidase inhibitor complexes adapted from (Gonçalves-Almeida, et. al., 2012). Clan (related tertiary structures) and family (related sequences) refer to MEROPS classification (Rawlings et. al., 2018). Class and fold account for SCOP structural classification (Murzin et. al., 1995).

| | ENZYME | | | INHIBITOR | | |
|---|---|---|---|---|---|---|
| **PDB** | **NAME SPECIE** | **CLAN FAMILY** | **CLASS FOLD** | **NAME SPECIE** | **CLAN FAMILY** | **CLASS FOLD** |
| 1ACB | α-Chymotrypsin *Bos taurus* | PA S1 | All-beta trypsin-like | Eglin C *Hirudo medicinalis* | IG I13 | Alpha and Beta CI-2 family |
| 1TEC | Thermitase *Thermoactinomyces vulgaris* | SB S8 | Alpha and Beta subtilisin-like | Eglin C *Hirudo medicinalis* | IG I13 | Alpha and Beta CI-2 family |
| 1CSE | Subtilisin Carlsberg *Bacillus subtilis* | SB S8 | Alpha and Beta subtilisin-like | Eglin C *Hirudo medicinalis* | IG I13 | Alpha and Beta CI-2 family |
| 1MEE | Mesentericopeptidase *Bacillus mesentericus* | SB S8 | Alpha and Beta subtilisin-like | Eglin C *Hirudo medicinalis* | IG I13 | Alpha and Beta CI-2 family |
| 1SBN | Subtilisin BPN *Bacillus amyloliquefaciens* | SB S8 | Alpha and Beta subtilisin-like | Eglin C L45R *Hirudo medicinalis* | IG I13 | Alpha and Beta CI-2 family |
| 1PPF | Leukocyte Elastase *Homo sapiens* | PA S1 | All-beta trypsin-like | Turkey Ovomucoid 3 *Meleagris gallopavo* | IA I01 | Small Protein/ Kazal-type |
| 1CHO | α-Chymotrypsin *Bos taurus* | PA S1 | All-beta trypsin-like | Turkey Ovomucoid 3 *Meleagris gallopavo* | IA I01 | Small Protein/ Kazal-type |
| 3SGB | SGT *Streptomyces griseus* | PA S1 | All-beta trypsin-like | Turkey Ovomucoid 3 *Meleagris gallopavo* | IA I01 | Small Protein/ Kazal-type |
| 1R0R | Subtilisin Carlsberg *Bacillus subtilis* | SB S8 | Alpha and Beta subtilisin-like | Turkey Ovomucoid 3 *Meleagris gallopavo* | IA I01 | Small Protein/ Kazal-type |

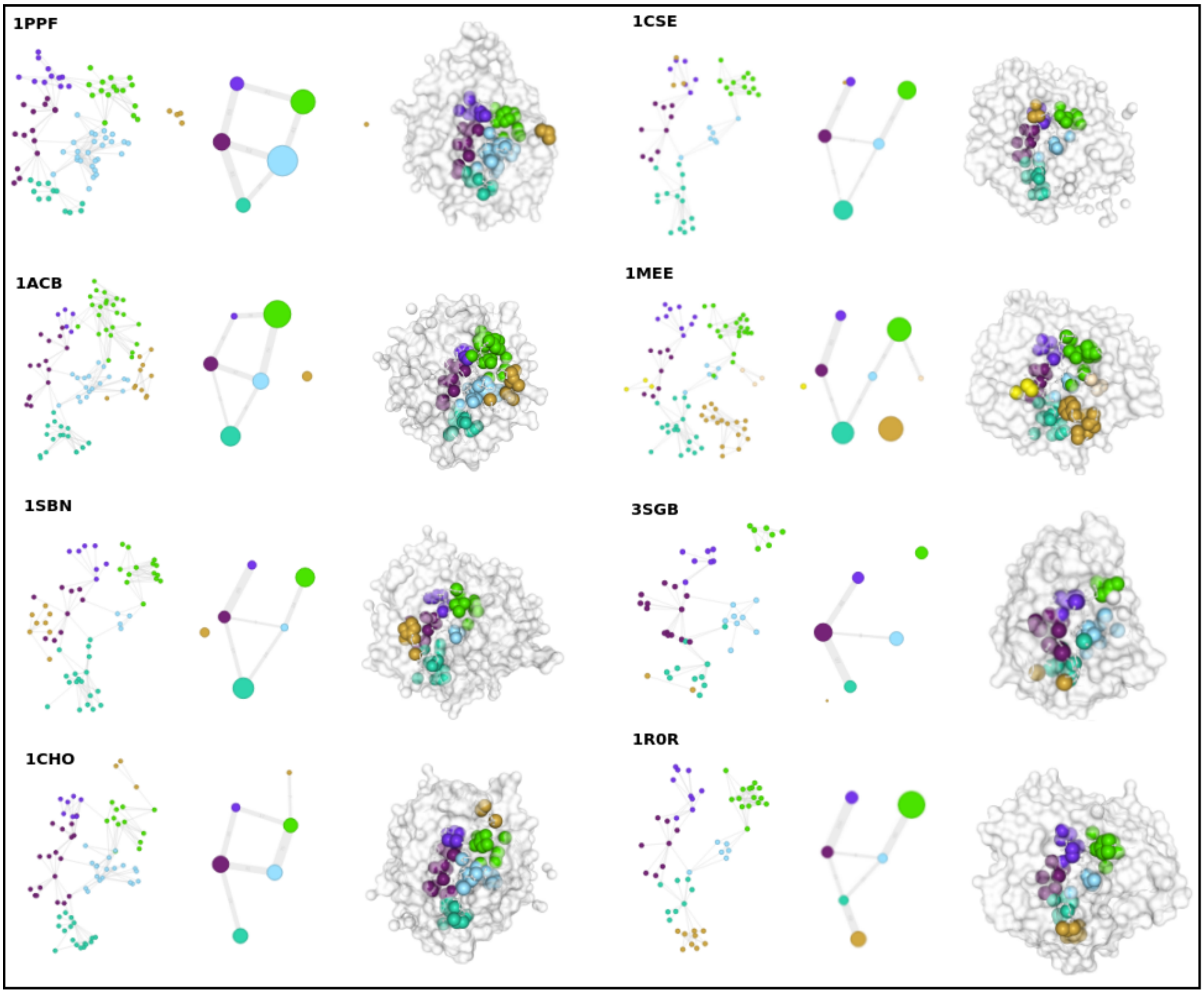


**Fig. S2.** Other higher graph alignments from the peptidase inhibitor dataset having 1PPF as a reference, showing lower-level graphs, higher-level graphs, and rendered PDB with surface and spacefill representations. This was automatically generated from the Topos algorithm and manually adjusted for better visualization. Graphs are only from nonpolar-nonpolar interactions, with the default internal Topos parameters: k = 4, n = 3, m = 8.

**Table S3.** Statistic tests for node size distribution comparisons (p-values)*

| Test | Type | Null hypothesis | Red-hot | Hot | Warm |
|---|---|---|---|---|---|
| Kolmogorov-Smirnov test | nonparametric | sample data come from the same distribution | 0.012 | 0.019 | 0.32 |
| Student’s t-test | parametric | sample data have the same means | 0.0033 | 0.0021 | 0.11 |
| Mann–Whitney–Wilcoxon test | nonparametric | sample data come from the same distribution | 0.0066 | 0.0038 | 0.10 |

* Calculated with R package - version 3.4.4 (R Core Team, 2018).

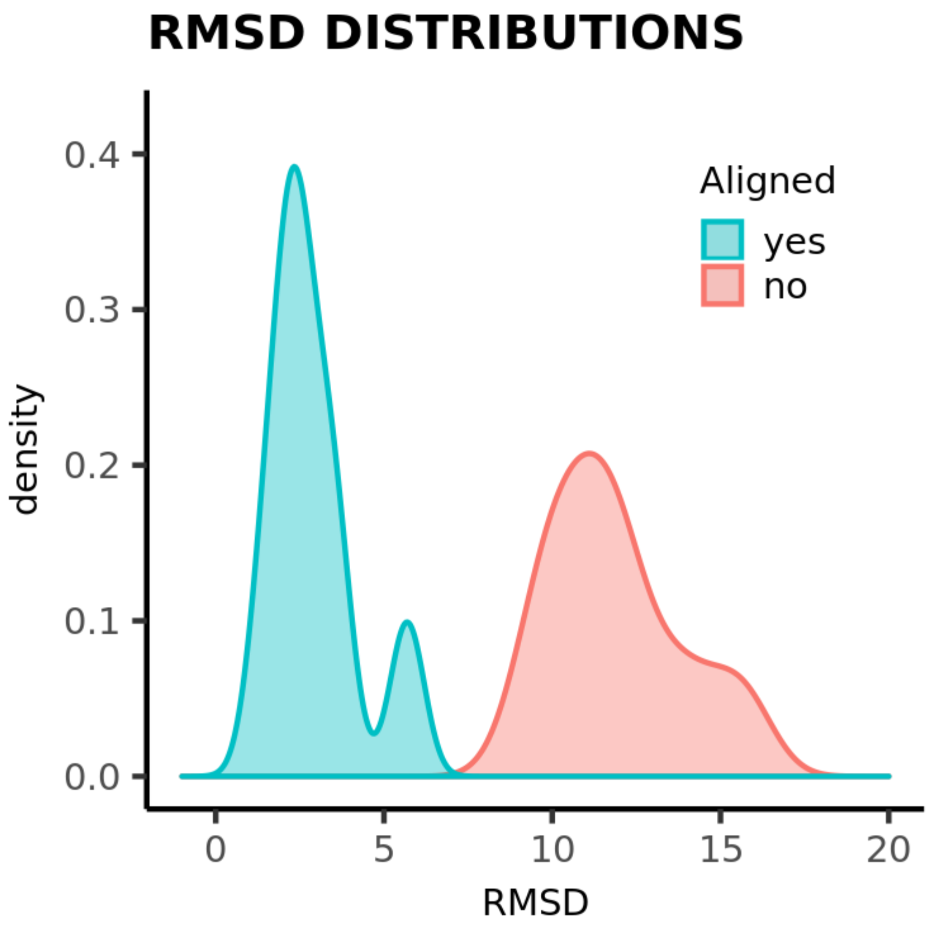


**Fig. S3.** RMSD (Root Mean Square Deviation) distributions for nodes position in aligned and non-aligned higher graphs of serine-peptidase inhibitor data set. PDB ids from Table S2 were aligned against 1PPF. RMSD averages are 2.9 Å and 11.9 Å, respectively.

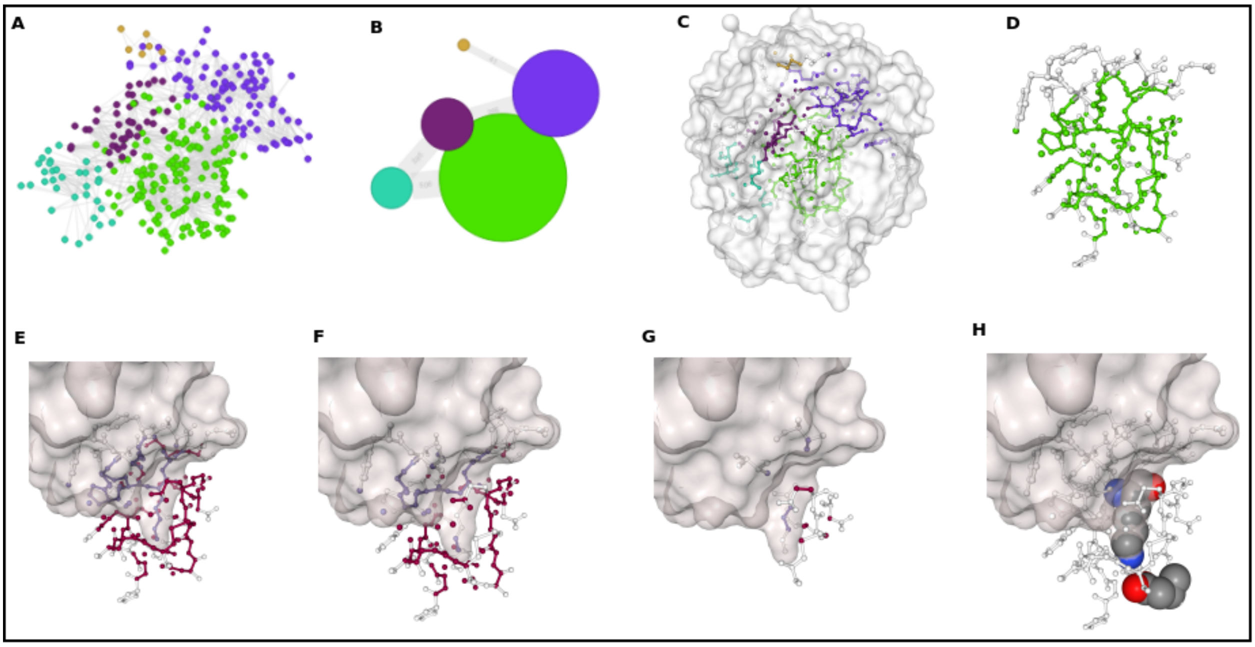

**Fig. S4.** Spot analysis of 2PTC, a bovine trypsin in complex with BPTI inhibitor. It has the biggest ΔΔG of binding (10 kcal.mol$^{-1}$) in the alanine scanning data set (Xia et al., 2010). (**A**) Lower-level graph with 5 partitions (colored in order to discriminate them), comprising ALL type of contacts (polar and nonpolar). (**B**) Higher-level graph representation. The mutation target in BPTI chain (K15) by alanine is located in the largest cluster (light-green) (**C**) Lower-level graph in the context of trypsin with surface (**D**) In higher graph, if user clicks on light-green node, this cluster can be detached from others. (**E**) Detached cluster colored by chain (trypsin in red, BPTI in blue) and with inhibitor surface representation. It is possible to see better now that this cluster has captured the trypsin catalytic pocket. (**F**) Focus on atoms with polar-polar interactions. (**G**) Focus on atoms with nonpolar-nonpolar interactions. It is noted that the pocket is essentially polar. (**H**) Two important residues of interface in salt bridge: BPTI-LYS15 and Trypsin-ASP189, highlighting the propensity of trypsins to bind positively charged residues into catalytic pocket. All of this analysis can be done simply with mouse, without any script or command-line.

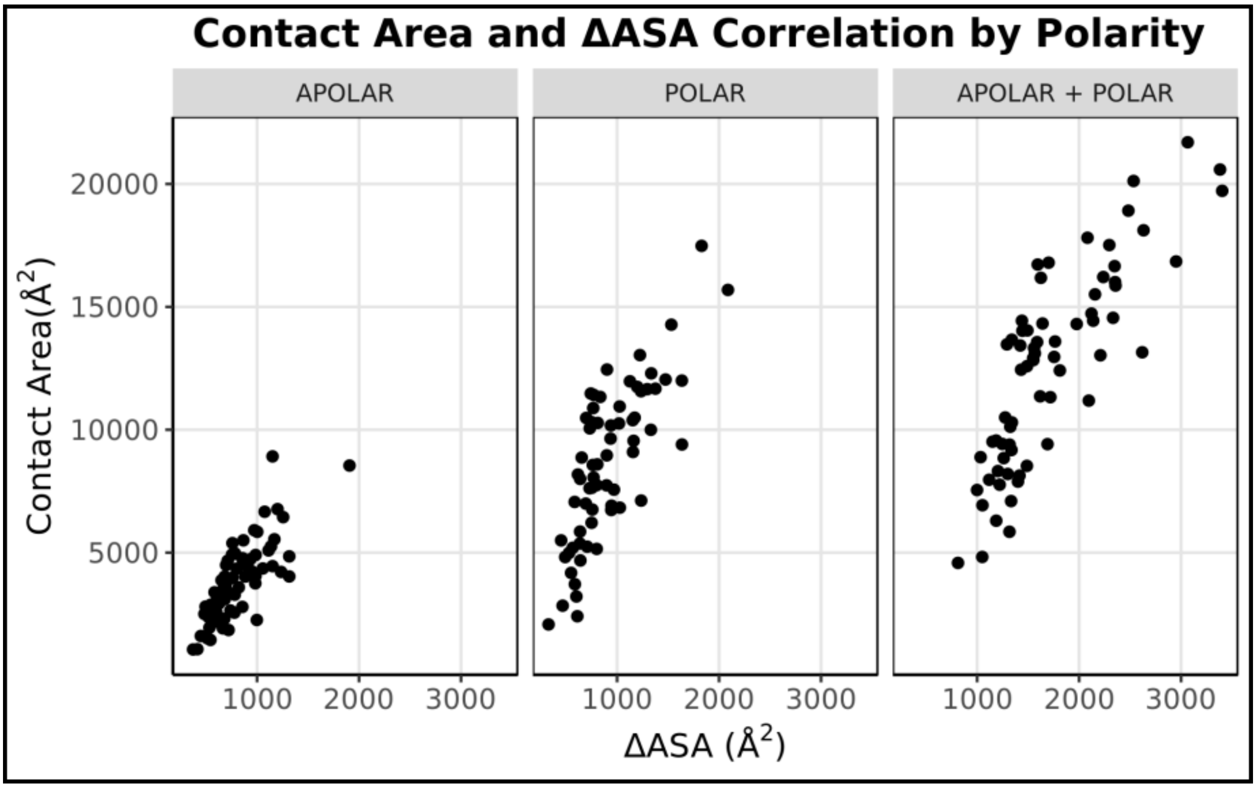


**Fig. S5.** Contact areas and ΔASA calculations from a sample of 68 chain-chain complexes from Affinity Database 2.0 (Kastritis et. al., 2011). ASAs were calculated using FreeSASA (Mitternacht, 2016) from vanddraabe R package (Esposito, 2017). This sample comprises only binary complexes considered to have a rigid-body binding, i.e., those having I-RMDS ≤ 1.5. I-RMDS is defined as a measure of the structural displacement when bound and unbound chains are compared (Hwang et. al., 2010). Pearson correlation was 0.79, 0.76 and 0.83 for apolar, polar, and both, respectively.

### *3) How to Generate the Figures Used in this Paper*

#### Figures 1A and S2:

It is important to emphasize again that the current version of GAPIN does not do mass alignments of PDBs, only pairwise alignments with two PDBs at a time.

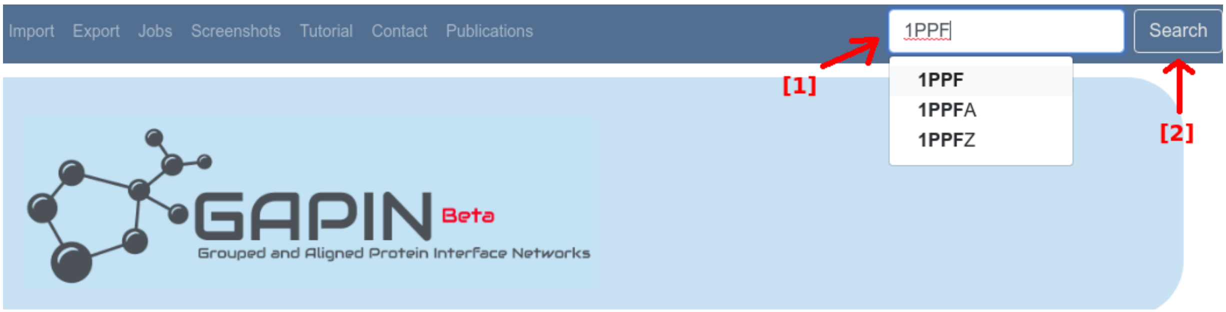


**STEP 1:** [1] enter or choose 1PPF, then [2] click on "Search"

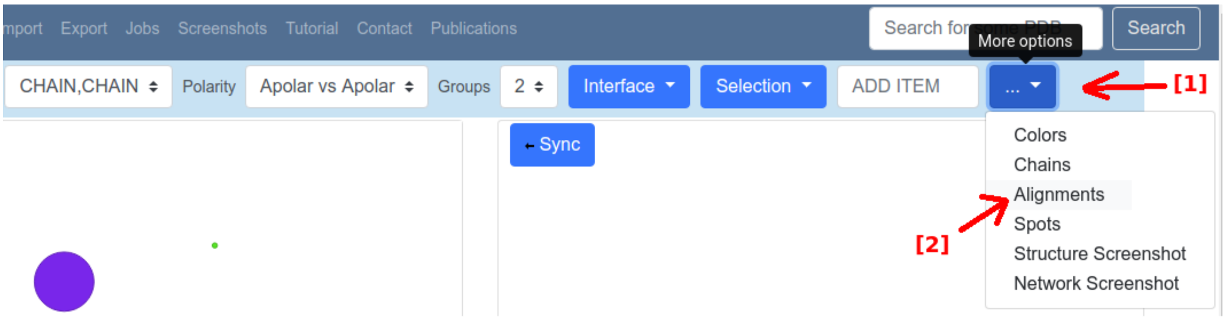


**STEP 2:** [1] click on "More option", then [2] choose "Alignments"

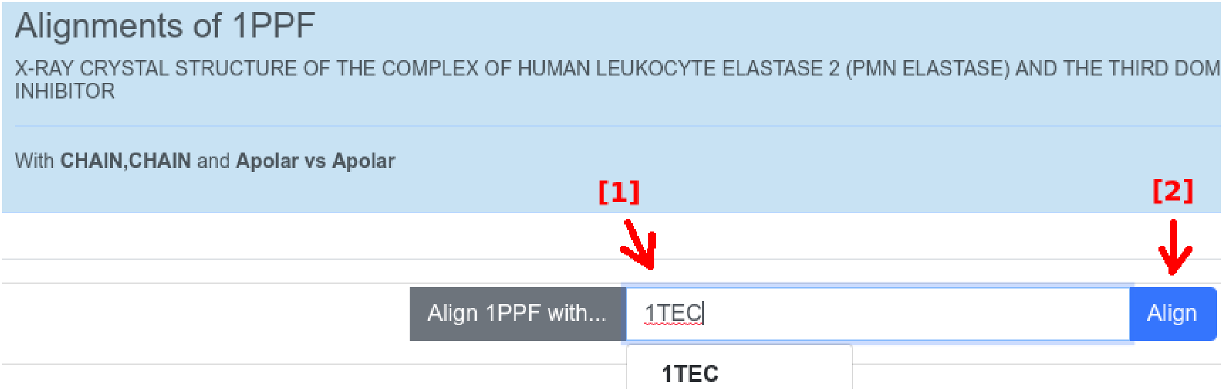


**STEP 3:** [1] enter or choose 1TEC, then [2] click on "Align"

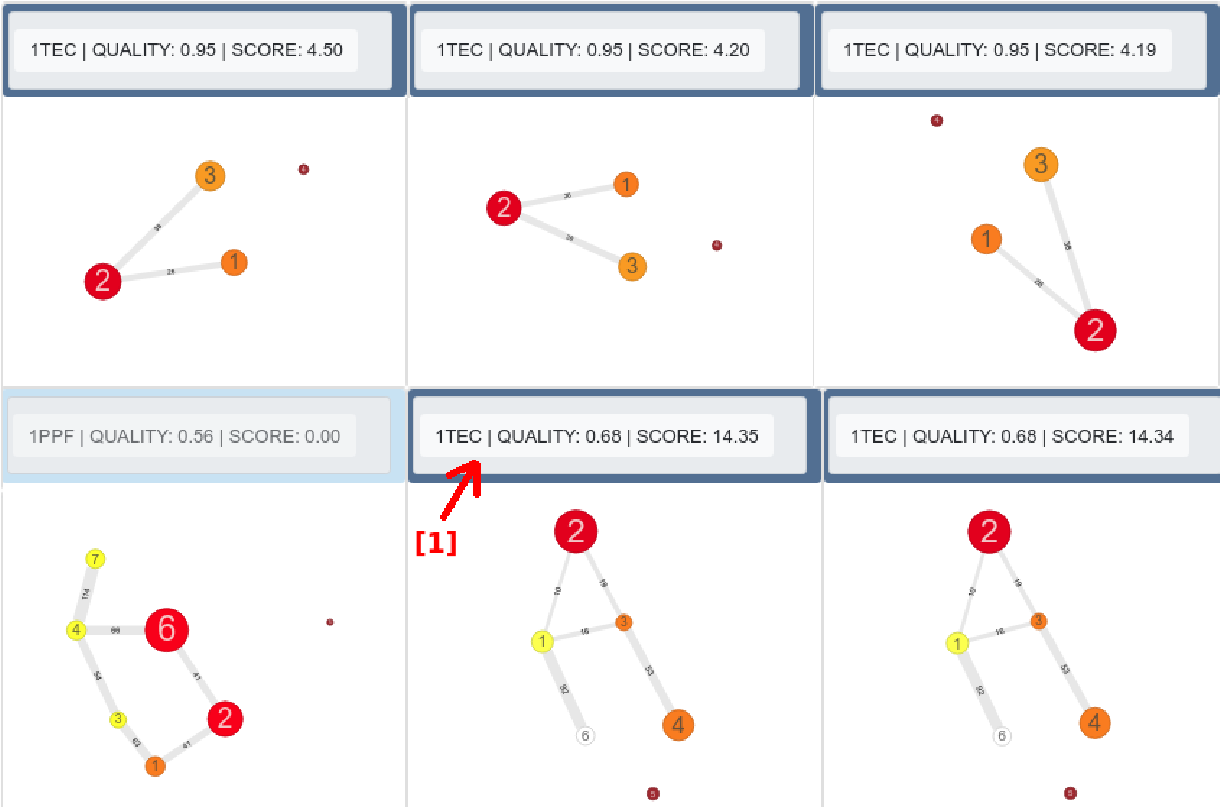


**STEP 4:** [1] choose appropriate quality and score.

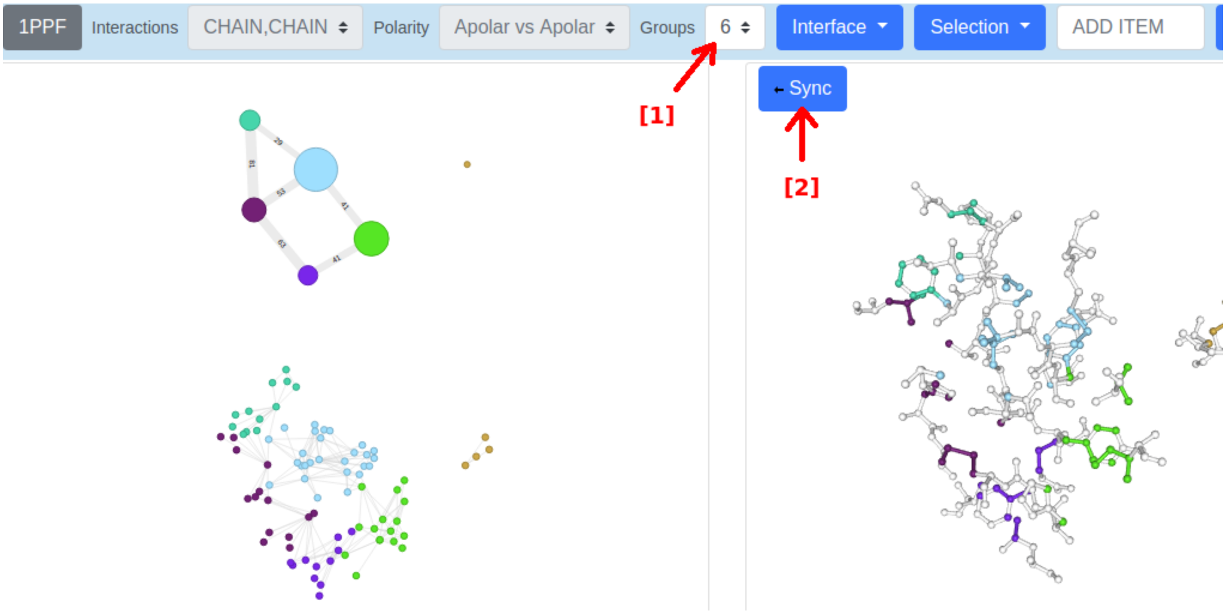


**STEP 5:** [1] you can change the number of groups without interfering with the final alignment. [2] Click on “Sync” to adjust rendered structure with graphs.

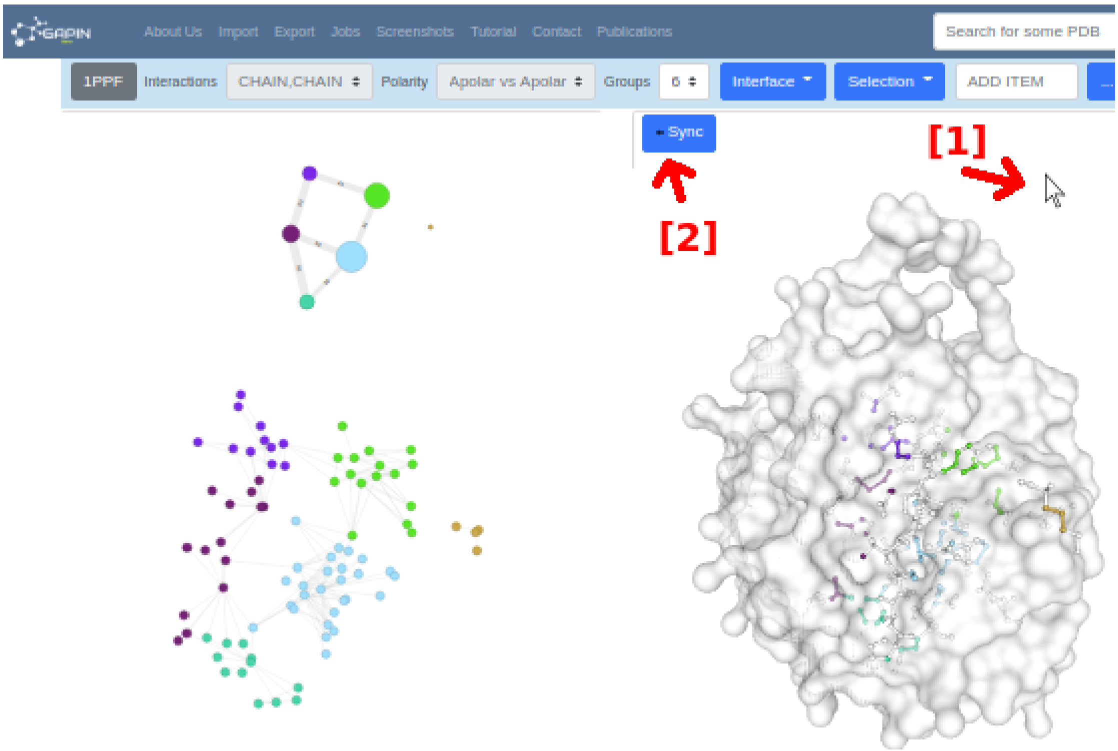


**STEP 6:** [1] as the plane of the interface will be pointing into the screen, with the mouse make the appropriate structure rotations (180° in Y and 180° in Z) to adjust it as shown. [2] Click on “Sync”. Do the same with 1TEC.

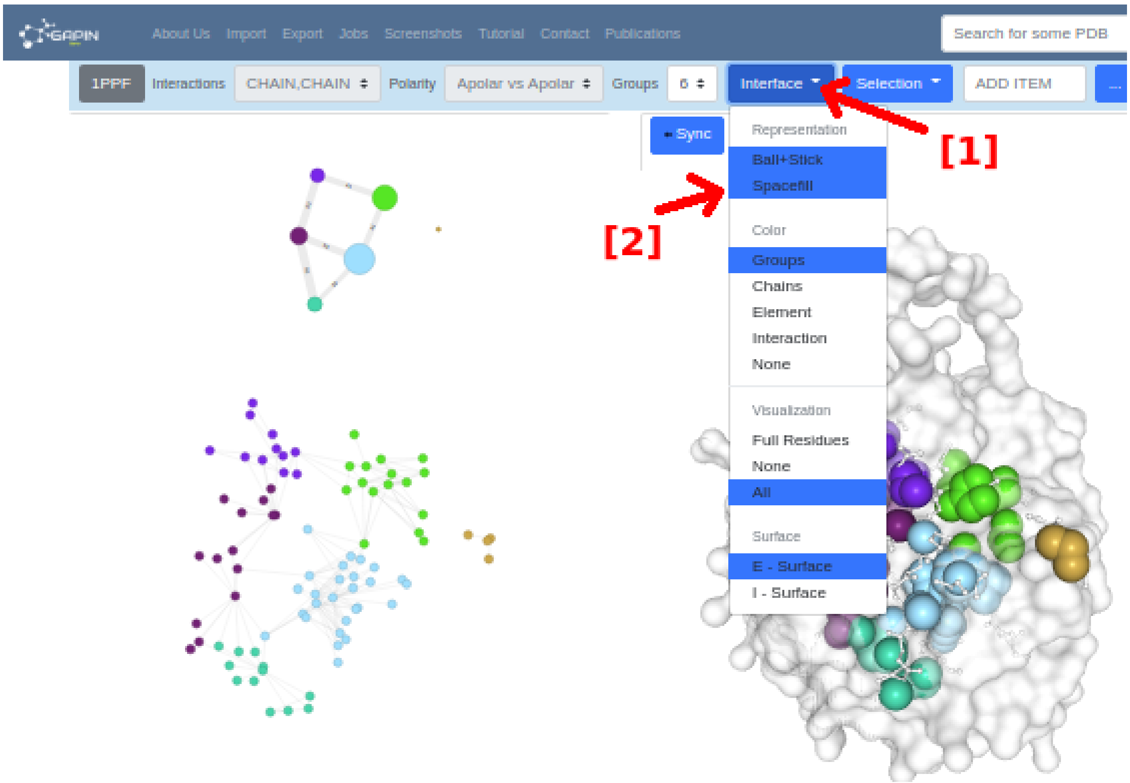


**STEP 7:** [1] click on “Interface”, then [2] on “Spacefill”. Do the same with 1TEC.

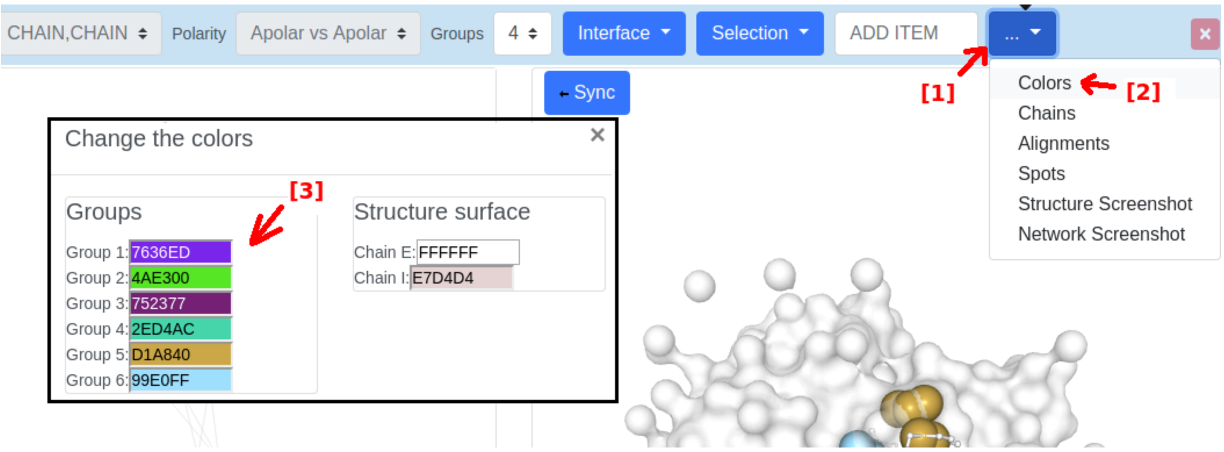


**STEP 8:** [1] in 1TEC, click "More options", then [2] "Colors", then [3] adjust with the appropriate colors to synchronize with 1PPF colors.

To generate the alignments shown in figure S2, change the PDB 1TEC for each PDB id listed there.

#### Figure S4:

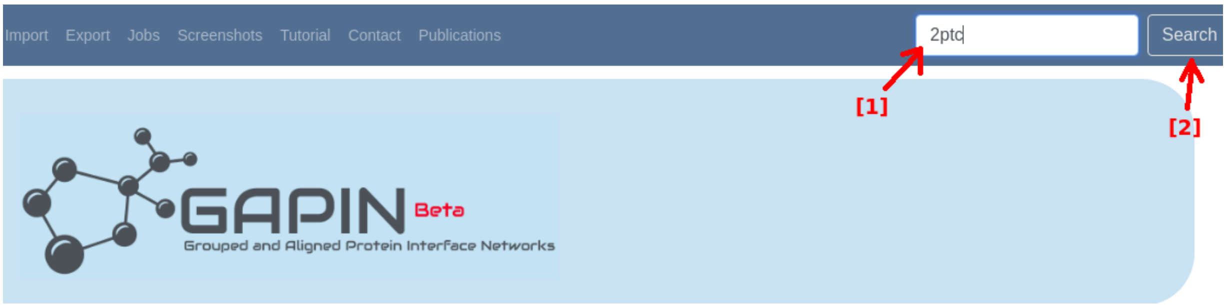


**STEP 1:** [1] enter or choose 2PTC, then [2] click on "Search"

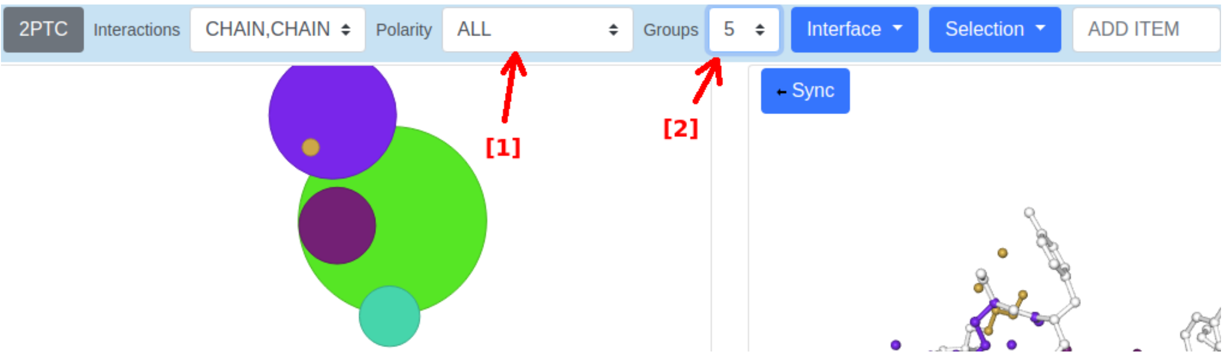


**STEP 2:** [1] click on "Polarity" and choose "ALL", then [2] click on "Groups" and choose "5".

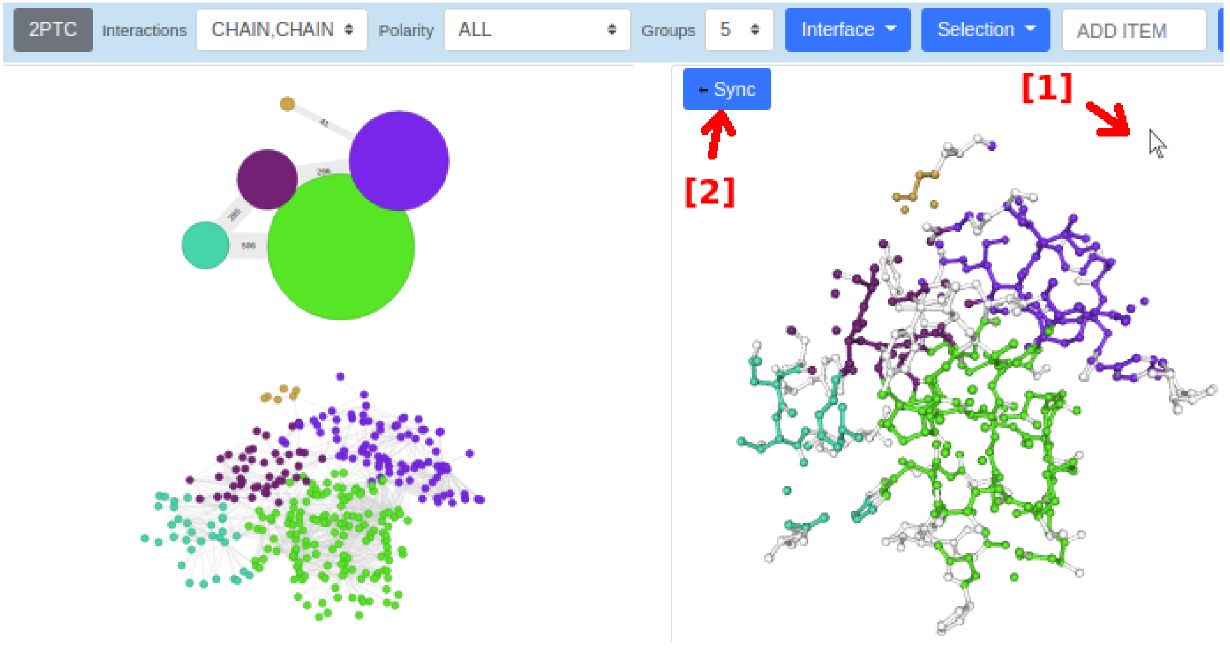


**STEP 3:** [1] with the mouse make the appropriate structure rotations to adjust it as shown, then [2] click on “Sync”.

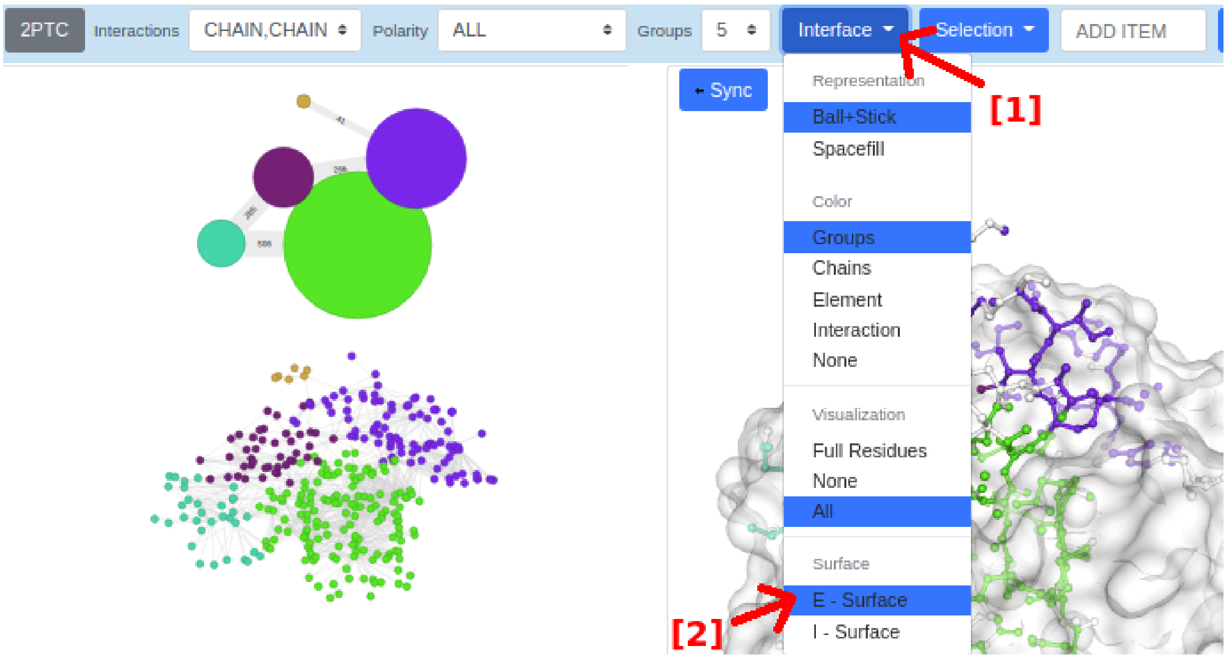


**STEP 4:** [1] click on “Interface”, then “E = Surface” (enzyme side).

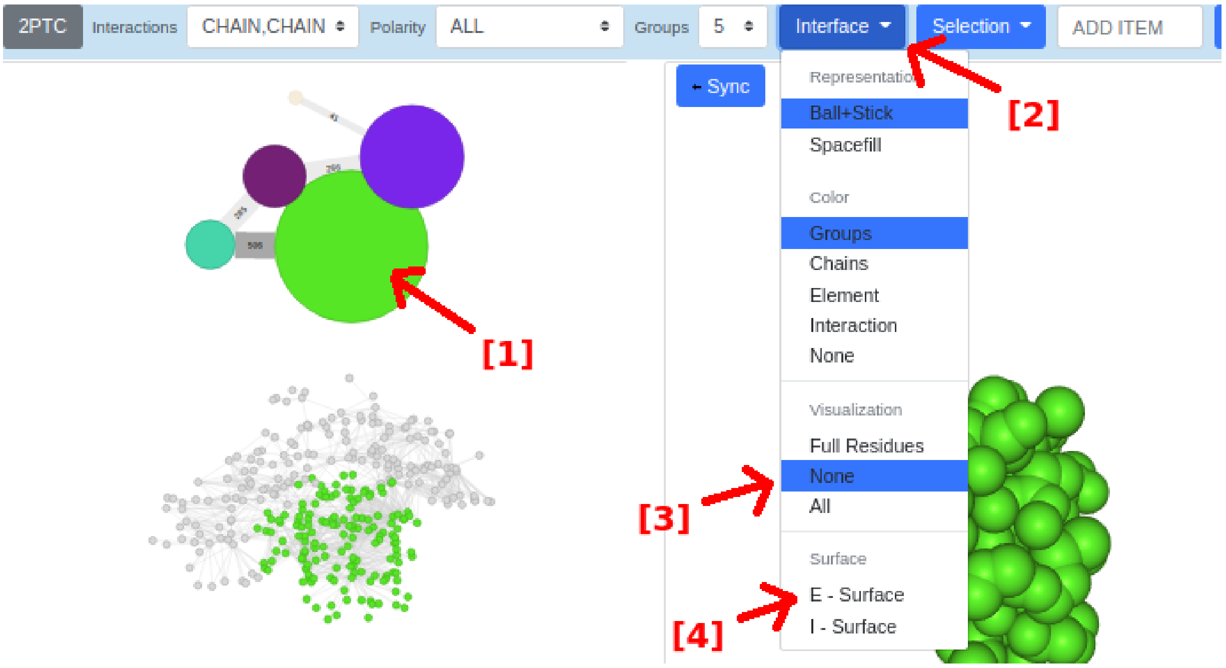


**STEP 5:** [1] click on the green and biggest higher graph node. [2] Click on “Interface”, then [3] “None” ( to isolated this node), then [4] “E-Surface” (to disable it).

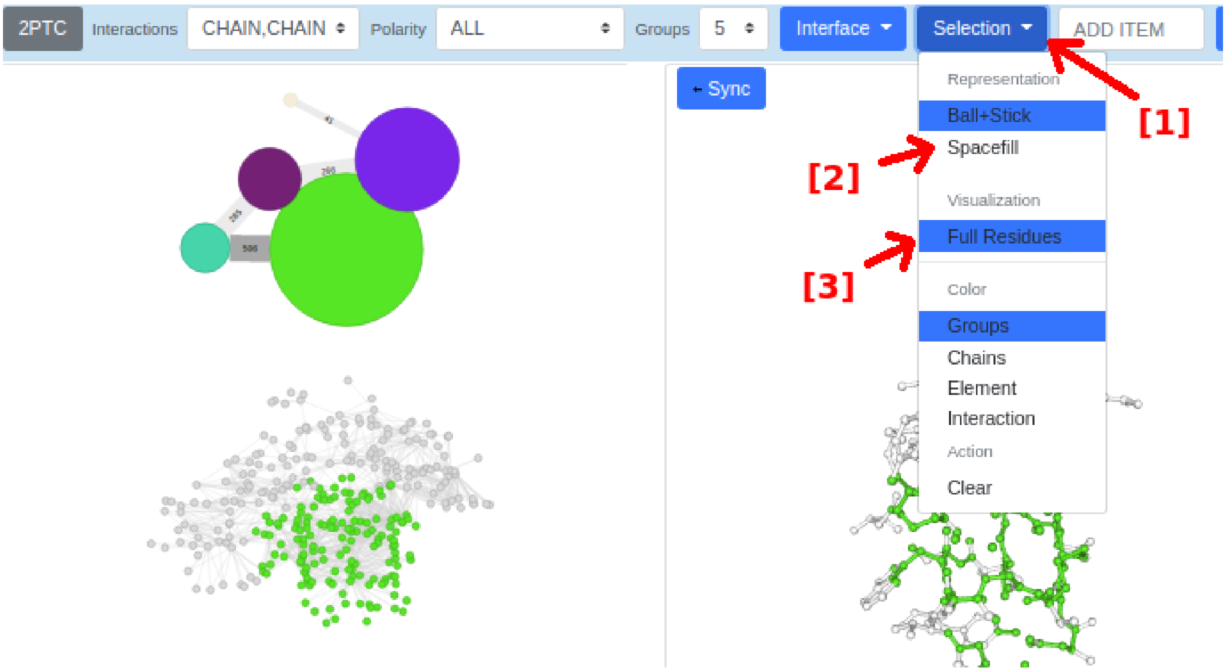


**STEP 6:** [1] click on “Selection”, then [2] “Spacefill” (to disable it), then “Full Residues” (to show also all residues atoms from this selection).

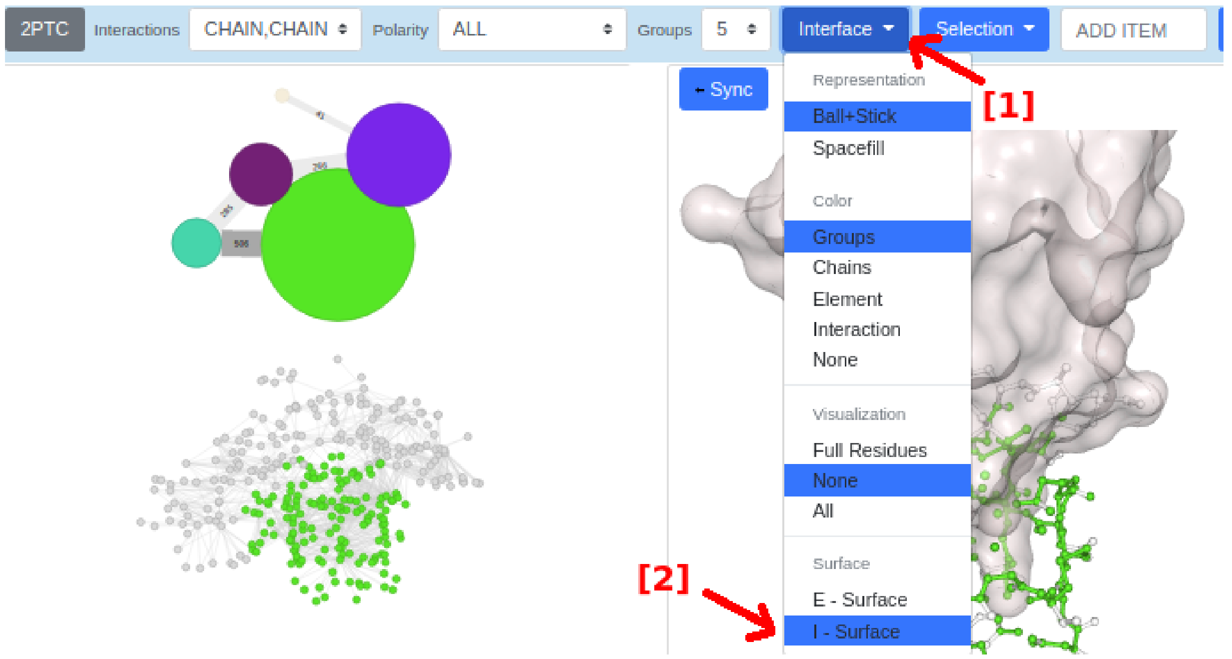


**STEP 7:** [1] click on “Interface”, then “I-Surface” to highlight now the inhibitor side.

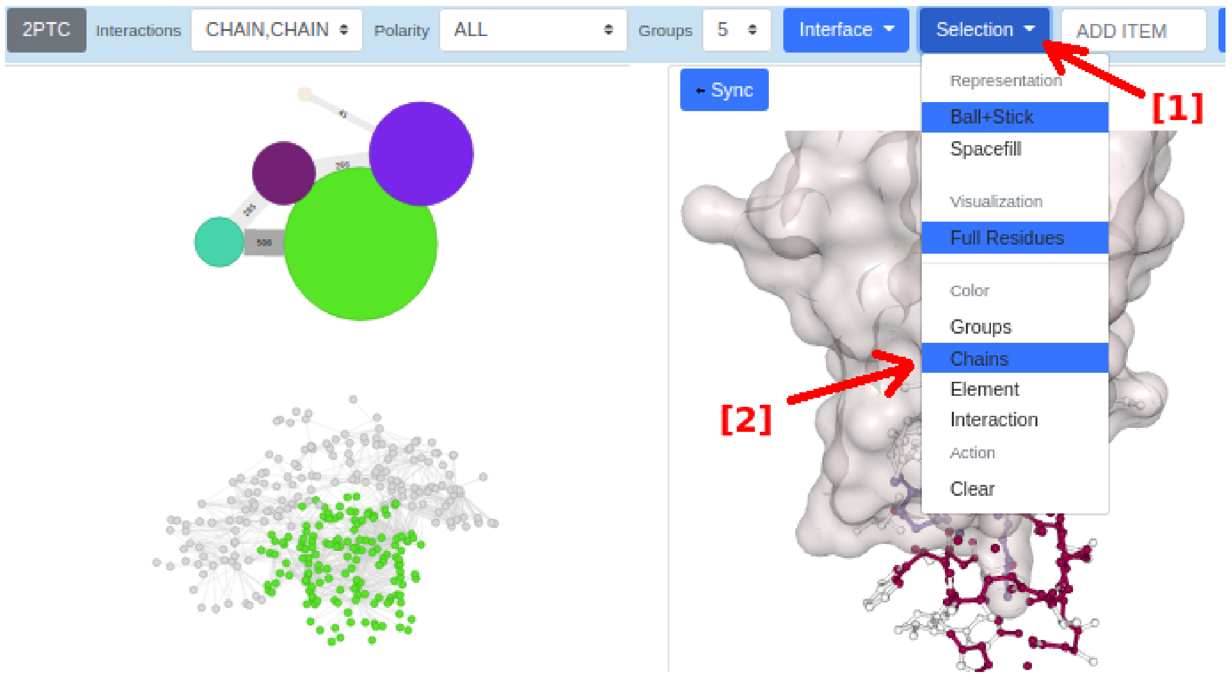


**STEP 8:** [1] click on “Selection”, then [2] “Chains”, to highlight colors by chains.

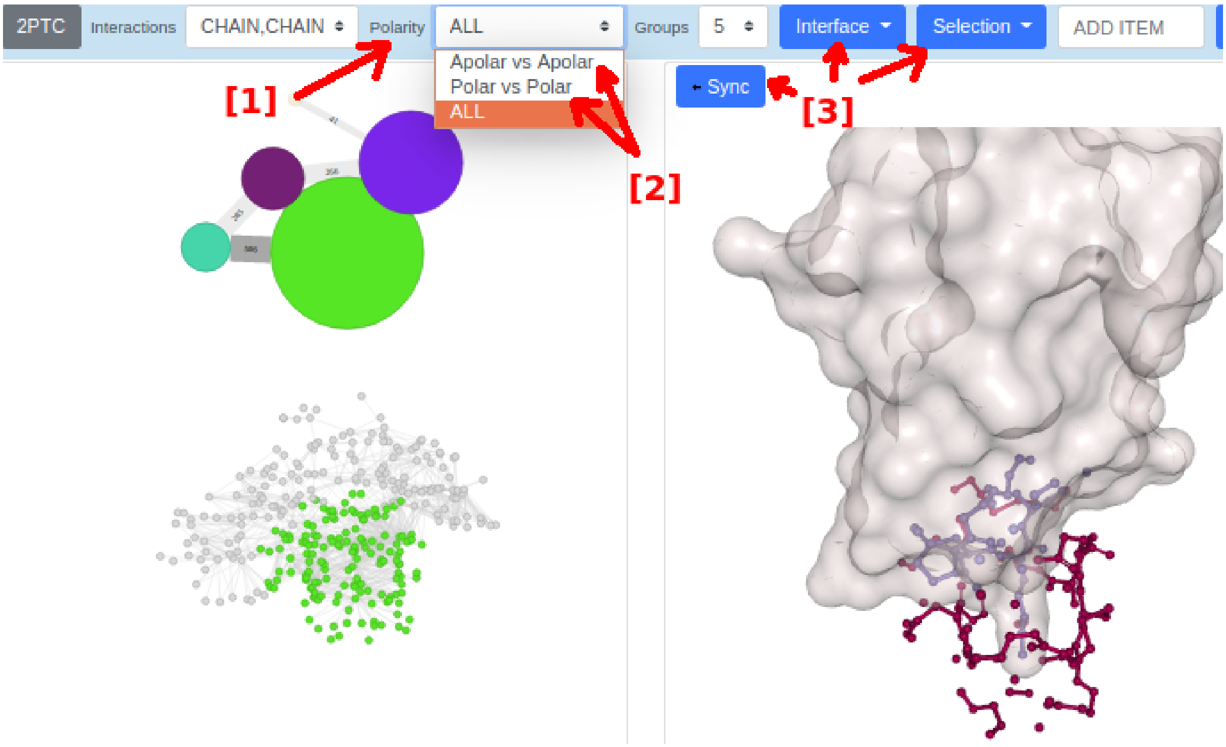


**STEP 9:** [1] click on "Polarity", then [2] change to "Apolar vs Apolar" or "Polar vs Polar" to focus on these types of contacts, then [3] make appropriate adjustments as in previous steps.

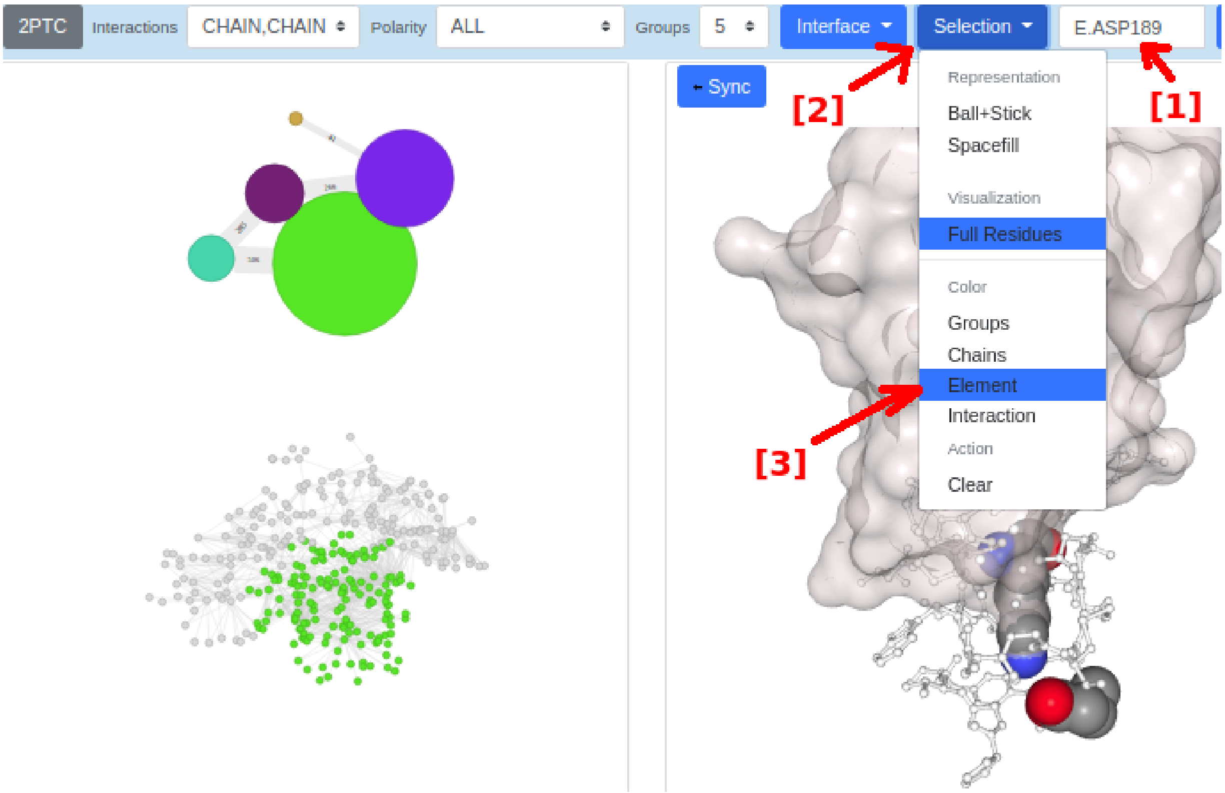


**STEP 10:** [1] to highlight specific atoms or residues (such as E.LYS15 and E.ASP189), use the autocomplete input box next to "Selection", then [2] click on "Selection", then [3] "Element" to color by atoms.

#### Figure S1:

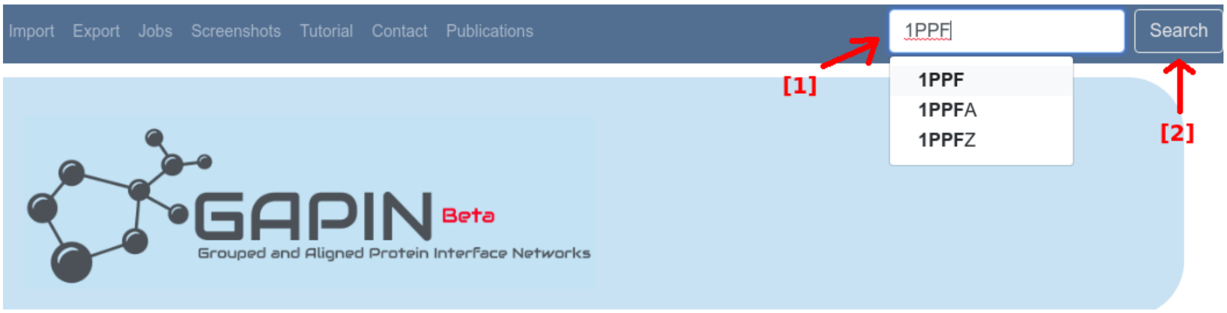


**STEP 1:** [1] enter or choose 1PPF, then [2] click on "Search"

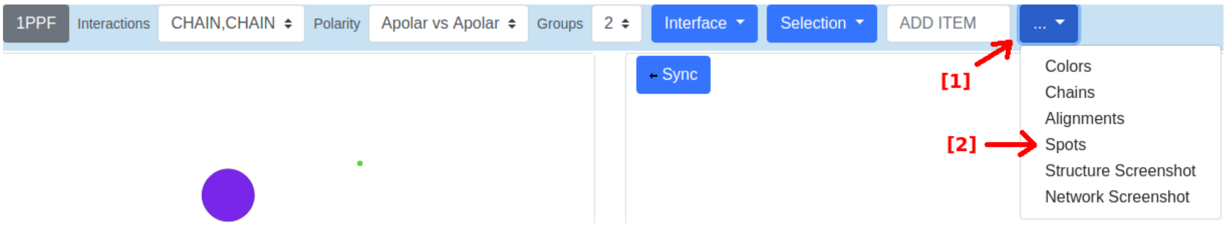


**STEP 2:** [1] click on "More option", then [2] choose "Spots"

### *4) Raw Data Used for Plots and Statistical Analysis*

See the file: GAPIN_supporting_raw_data_files-v1.zip

### *5) References*


da Silveira, C. H., Pires, D. E. V, Minardi, R. C., Ribeiro, C., Veloso, C. J. M., Lopes, J. C. D., … Santoro, M. M. (2009). Protein cutoff scanning: A comparative analysis of cutoff dependent and cutoff free methods for prospecting contacts in proteins. Proteins, 74(3), 727–43.

Esposito, E. (2017, April). Vanddraabe: Identification and statistical analysis of structurally conserved waters via R. In Abstracts Of Papers Of The American Chemical Society (Vol. 253). 1155 16th St, Nw, Washington, Dc 20036 Usa: Amer Chemical Soc.

Gonçalves-Almeida, V. M., Pires, D. E. V, de Melo-Minardi, R. C., da Silveira, C. H., Meira, W., & Santoro, M. M. (2012). HydroPaCe: understanding and predicting cross-inhibition in serine proteases through hydrophobic patch centroids. Bioinformatics (Oxford, England), 28(3), 342–9.

Hubbard,S.J. and Thornton,J.M. (1993) Naccess. Computer Program, Department of Biochemistry and Molecular Biology. University College, London.

Hwang, H., Vreven, T., Janin, J., & Weng, Z. (2010). Protein-protein docking benchmark version 4.0. Proteins: Structure, Function and Bioinformatics, 78(15), 3111–3114.

Kastritis, P. L., Moal, I. H., Hwang, H., Weng, Z., Bates, P. a., Bonvin, A. M. J. J., & Janin, J. (2011). A structure-based benchmark for protein-protein binding affinity. Protein Science, 20(3), 482-91.

Kelly, PA. (2019). Mechanics Lecture Notes Part III: Foundations of Continuum Mechanics. URL:http://homepages.engineering.auckland.ac.nz/~pkel015/SolidMechanicsBooks/Part_III/Chapter_1_Vectors_Tensors/Vectors_Tensors_05_Coordinate_Transformation_Vectors.pdf (11 Fev 2019, date last accessed)

Lancichinetti, A., & Fortunato, S. (2009). Community detection algorithms: A comparative analysis. Physical Review E - Statistical, Nonlinear, and Soft Matter Physics, 80(5), 1–11.

Lee, B., & Richards, F. M. (1971). The interpretation of protein structures: Estimation of static accessibility. Journal of Molecular Biology, 55(3).

Mitternacht, S. (2016). FreeSASA: An open source C library for solvent accessible surface area calculations. F1000Research, 5(189), 1–10.

Murzin, A. G., Brenner, S. E., Hubbard, T., & Chothia, C. (1995). SCOP: A structural classification of proteins database for the investigation of sequences and structures. Journal of Molecular Biology, 247(4), 536–540.

Newman, M. E. J. (2004). Detecting community structure in networks. European Physical Journal B, 38(2), 321–330.

Newman, M. E. J., & Girvan, M. (2004). Finding and evaluating community structure in social networks. Physical Review E, 69(2), 1–15.

Rawlings, N. D., Barrett, A. J., Thomas, P. D., Huang, X., Bateman, A., & Finn, R. D. (2018). The MEROPS database of proteolytic enzymes, their substrates and inhibitors in 2017 and a comparison with peptidases in the PANTHER database. Nucleic Acids Research, 46, D624-D632

Rousseeuw, P. J. (1987). Silhouettes: A graphical aid to the interpretation and validation of cluster analysis. Journal of Computational and Applied Mathematics, 20(C), 53–65.

Sobolev, V., Sorokine, A., Prilusky, J., Abola, E. E., & Edelman, M. (1999). Automated analysis of interatomic contacts in proteins. Bioinformatics, 15(4), 327–332.

White, S. H., & Wimley, W. C. (1999). Membrane protein folding and stability: physical principles. Annual Review of Biophysics and Biomolecular Structure, 28, 319–365.

Zaki, M.J., Meira Jr, W., (2014). Data mining and analysis: fundamental concepts and algorithms. New York : Cambridge University Press.

R Core Team (2018). R: A language and environment for statistical computing. R Foundation for Statistical Computing, Vienna, Austria. URL https://www.R-project.org/.

### A APPENDIX: Demonstration of BARS Equation

Let's $R_1$ and $R_2$ be the radii of two spheres $S_1$ and $S_2$, respectively, the first with a center in the origin:

$$x^2+y^2+z^2=R_1^2 \quad (1)$$

and the second with a center over z axis given by:

$$x^2+y^2+(z-d)^2=R_2^2 \quad (2)$$

where $d>0$ (distance from origin) e $R_1 \geq R_2$. Consider now each of the spherical caps generated by the intersection of the two spheres using the double integral for generic surfaces:

$$A=\iint_D \sqrt{f_x(x,y)^2+f_y(x,y)^2+1}\,dA \quad (3)$$

We need to find a region $D$ and the functions that describe the surfaces.

Joining (1) and (2), we have:

$$R_1^2-z^2+(z-d)^2=R_2^2 \quad (4)$$

With gives:

$$z=\frac{d^2+R_1^2-R_2^2}{2d} \quad (5)$$

Replacing $z$ in equation (1) gives the region of integration $D$:

$$x^2+y^2=R_1^2-\frac{d^2+R_1^2-R_2^2}{2d} \quad (6)$$

Now, isolating $z$ in equation (1) gives a function $z=f(x,y)$:

$$z=\sqrt{R_1^2-x^2-y^2} \quad (7)$$

With the following partial derivatives:

$$\frac{\partial f}{\partial x} = -\frac{1}{\left(R_1^2 - x^2 - y^2\right)^{1/2}} x$$
$$\frac{\partial f}{\partial y} = -\frac{1}{\left(R_1^2 - x^2 - y^2\right)^{1/2}} y \tag{8}$$

Replacing (8) in (3), we have:

$$A(S_1) = \iint_D \frac{R_1}{\sqrt{R_1 - x^2 - y^2}} dxdy \tag{9}$$

Rewriting (9) in polar coordinates:

$$A(S_1) = -\frac{R_1}{2} \int_0^{2\pi} \int_{R_1^2}^{M^2} \frac{1}{\sqrt{u}} dud\theta$$
$$where$$
$$M = \frac{d^2 + R_1^2 - R_2^2}{2d}, \; u = R_1^2 - r^2 \tag{10}$$

Solving (10) gives:

$$A(S_1) = 2\pi\left[R_1^2 - R_1 M\right] \tag{11}$$

Similarly:

$$A(S_2) = 2\pi\left[R_2^2 - R_2 M\right] \tag{12}$$

Doing the math by adding (11) and (12), we finally have:

$$A(R_1, R_2, d) = 2\pi\left(R_1^2 + R_2^2\right) - \pi\left(R_1 + R_2\right) d\left[1 + \left(\frac{R_1 - R_2}{d}\right)^2\right] \tag{13}$$
